# PRRGO: A Tool for Visualizing and Mapping Globally Expressed Genes in Public Gene Expression Omnibus RNA-Sequencing Studies to PageRank-scored Gene Ontology Terms

**DOI:** 10.1101/2024.01.21.576540

**Authors:** Luis E. Solano, Nicholas M. D’Sa, Nikolas Nikolaidis

## Abstract

We herein report PageRankeR Gene Ontology (PRRGO), a downloadable web application that can integrate differentially expressed gene (DEG) data from the gene expression omnibus (GEO) GEO2R web tool with the gene ontology (GO) database [1]. Unlike existing tools, PRRGO computes the PageRank for the entire GO network and can generate both interactive GO networks on the web interface and comma-separated values (CSV) files containing the DEG statistics categorized by GO term. These hierarchical and tabular GO-DEG data are especially conducive to hypothesis generation and overlap studies with the use of PageRank data, which can provide a metric of GO term centrality. We verified the tool for accuracy and reliability across nine independent heat shock (HS) studies for which the RNA-seq data was publicly available on GEO and found that the tool produced increasing concordance between study DEGs, GO terms, and select HS-specific GO terms.

## INTRODUCTION

The National Center for Biotechnology Information (NCBI) maintains gene expression omnibus (GEO), a staple repository of publicly accessible gene expression and functional genomics datasets [2]. GEO enables scientists with an internet connection to promote reproducibility in the biological research space by hosting a venue for depositing, sharing, and accessing gene expression data sourced from various model organisms generated by micro-array, sequencing, and high-throughput next-generation sequencing (NGS) technologies. The standardized format for data submission also facilitates the use of tools attempting to take advantage of the wealth of information.

For publicly available NGS gene expression studies, whether researchers choose to download FASTQ files, raw count matrices, or processed normalized count matrices, researchers must be knowledgeable regarding tools to calculate differentially expressed genes (DEGs). DESeq2 [3], edgeR [4], limma [5], and voom [6] are examples of heavily cited tools to achieve this important step in the analysis of gene expression data. A full characterization, review, and benchmarking of these tools is beyond the scope of this article and achieved more comprehensively in Seyednasrollah et al [7].

The GEO2R feature of the Gene Expression Omnibus (GEO) allows users to calculate differential gene expression with DESeq2 on NCBI servers [3]. In addition to outputting both visual and tabular formatted processed DEG data, this tool enables users to compute their own DEG data for any publicly available gene expression dataset hosted on GEO. Once a user has secured these DEGs, their corresponding fold change and p-values are frequently used as inputs into various tools for enrichment and pathway analysis steps. These tools often base their reference annotations upon the Gene Ontology (GO) [1, 8]. GO contains information about biological processes (BP), molecular functions (MF), and cellular components (CC) organized in a hierarchical network graph.

### The PageRank Algorithm

The PageRankeR Gene Ontology (PRRGO) visualization and exploration tool utilizes the PageRank algorithm in a novel way for calculating the relative importance of GO terms. The PageRank algorithm was originally used to rank web pages [9, 10], but now its use spans many different fields. For any network graph with nodes and directed edges, PageRank assigns relative weights to each node based on the number of inbound links for each node. Each algorithm iteration simulates a web user that begins at a node chosen at random; the user randomly clicks on links represented by edges to arrive at different web pages represented by nodes [10]. PageRank then computes the percent chance of arriving at any given webpage [10]. Notably, PageRank has been applied to many biological contexts including identifying candidate genes [11], topologically expressed genes [12], protein function prediction [13], gene evaluation from microarray results [14], finding functional gene modules [15, 16], entity linking [17], prioritizing transcriptional factors in gene regulatory networks [18], and semantic similarity for disease-target associations [19].

### GO Related Applications

Many previous software applications have enabled the user to annotate GO terms to genes. ClueGO [20] is a Cytoscape [21] plug-in that is capable of importing and performing enrichment analyses given a list of DEGs for both the GO and Kyoto Encyclopedia of Genes and Genomes (KEGG) databases [22]. ClueGO performs network visualization with nodes colored based on GO enrichment results and clustering of functionally related groups of terms [20]. GOnet is a web application that performs GO annotation or GO term enrichment analysis for a given list of genes with their contrast values [23]. The results of these analyses can be viewed and exported in both an interactive graphical format and as comma-separate values (CSV) and text (.txt) files. NaviGO is a GO visualization and enrichment application that depicts the hierarchy of the GO network when used in conjunction with its linked GO Visualizer [24]. Similar to ClueGO, NaviGO also computes enrichment scores for GO terms and color GO term nodes in a network based on these scores. WEGO is a web application that produces descriptive statistics for GO terms given a list of enriched genes [25]. WEGO produces bar graphs showing the number and percentage of genes annotated to a specific GO term and can perform GO enrichment analysis. AEGIS is a downloadable application that generates interactive focus and context graphs that provide a novel way of exploring GO terms for DEGs [26]. It also allows the user to focus on certain gene sets and perform power analysis. AmiGO is a web application that allows searching for GO terms based on keywords and viewing the genes annotated to those keywords [27]. It also provides the ability to visualize and export the annotations and GO network information.

We see three key deficiencies present in the current GO software ecosystem that we hope to address with PRRGO:

1. Current tools do not allow importing complete DEG data (i.e. including multiple fields like log2 fold change and p-values) and annotating DEG data with relevant GO terms. At most, only the genes and contrast values can be parsed and exported in relevant data files in current tools.
2. The relative importance and seniority of individual GO terms is often lost when GO-DEG annotation files are produced. While the interactive network graphs do show the hierarchy, a quantitative metric in data files would allow comparison and prioritizing of such terms for further exploration.
3. The data files that can be exported from current tools vary in their usefulness to downstream analysis and visualization. The current files that can be exported allow contrast values to be connected to DEGs and GO terms, but all three cannot be viewed simultaneously without further data processing. Data should be easily comparable across studies, DEGs, and GO terms for analysis and visualization.

In developing PRRGO, our aim is to bridge these gaps. We offer researchers a versatile and user-friendly tool that not only overcomes existing deficiencies, but also introduces novel functionalities, notably the application of the PageRank algorithm for quantitatively assessing the centrality of GO terms. The subsequent sections detail the features, validation, and application of PRRGO, showcasing its potential to enhance the landscape of GO-DEG analysis tools.

## METHODS

### PRRGO Software Architecture and Query Handling

The PRRGO software architecture (Figure 1) integrates DEG and GO data. These data are displayed in both graphical and tabular formats. This network visualization and differentially expressed gene mapper is hosted on a Django web server that can be run locally on Linux, Windows, and Mac operating systems. PRRGO queries to the server comprising a keyword, the number of terms to filter by PageRank score, and file paths to common DEG export files, are transmitted via HTTP POST.

**FIGURE 1.**
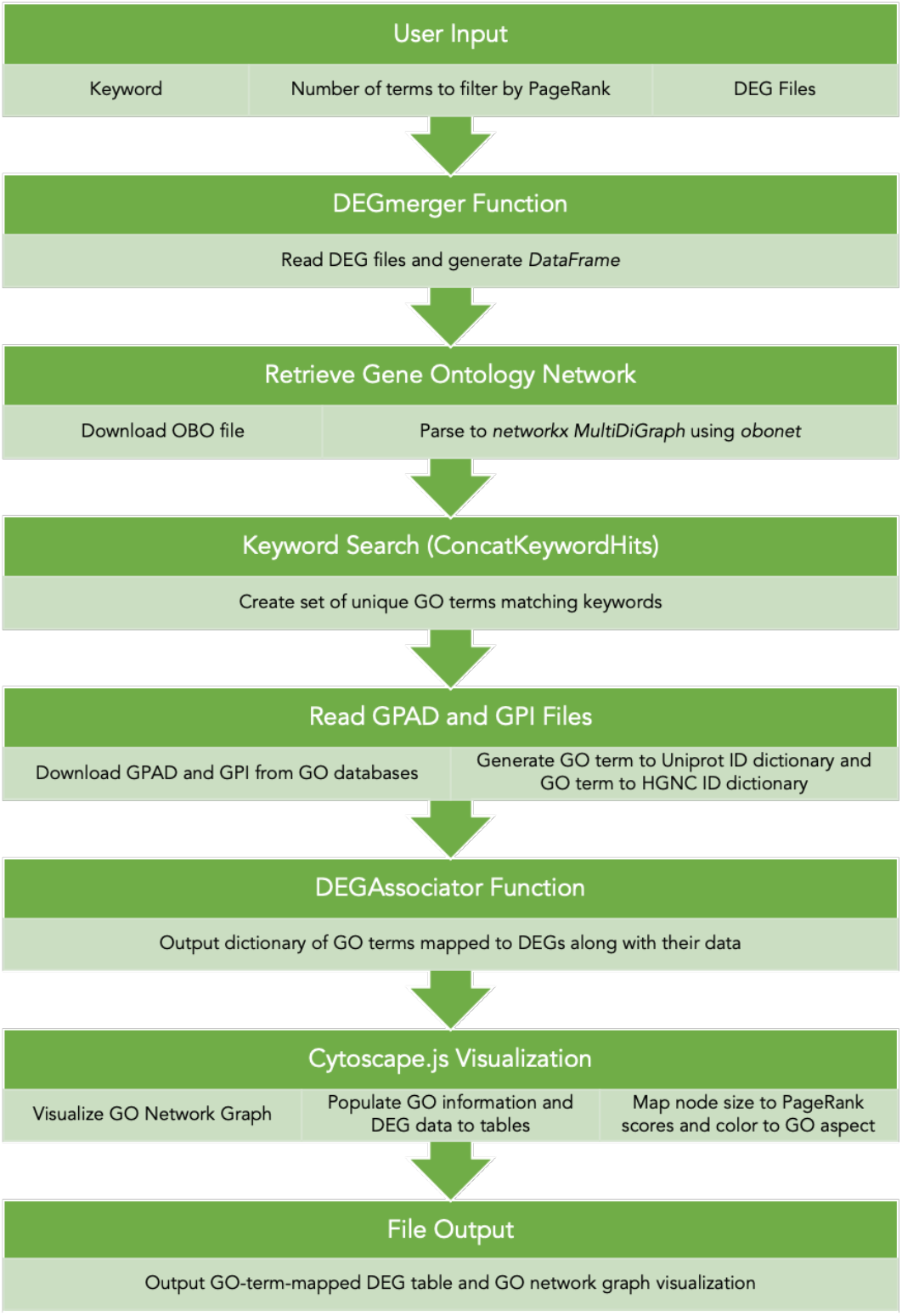
PRRGO ARCHITECTURE. PRRGO first accepts a keyword string and reads in GEO2R DEG exports from user-provided file paths; these are converted to a dataframe and file names appended. An OBO formatted GO network is made locally available, converted to a network graph, and a keyword search is executed against this network following the calculation of pagerank scores. Super-users will find various supporting functions based on relational GO annotations. Next GPAD and GPI files supply PRRGO with reference annotations and identifiers by which to map them within a dictionary created by the DEGassociator function. Lastly, Cytoscape.js provides an architecture for interactive visualization of the mapped relationships with an option to export critical information as a CSV file.

### DEG Data Integration and Gene Ontology Network Retrieval

The DEGmerger function processes user-supplied file paths to GEO2R-generated DEG export files. After determining the delimiter format, it constructs a dataframe from supplied DEG statistics and appends the filename of each respective DEG export file to corresponding columns enabling multiple DEG export file inputs. Next, the current Gene Ontology (GO) network is retrieved and locally stored as an OBO-formatted graph accessible via http://purl.obolibrary.org/obo/go.obo. The networkx package [28] converts the OBO-based GO graph into a network graph, and then computes PageRank scores for that entire network graph via the PageRank function [9, 10].

### Keyword Search and Filtering

PRRGO then executes a keyword search through all GO term names and descriptions creating a set of unique GO terms containing the specified keyword either in the GO term name or description. The KWNameQuery, KWDefQuery, and ConcatKeywordHits functions work together to process the user keyword input, cross referencing against GO term names and definitions, then collecting network node names that represent unique hits.

### Visualization with Cytoscape.js

After filtering by top n PageRank scores, Cytoscape.js [29] JSON format facilitates the association of GO terms with their GO name, aspect, definition, PageRank scores, and DEG relevant data. The data are then sent to the user’s web browser, where the Cytoscape.js [29] library visualizes the GO network. The visualized network includes size-mapped nodes based on PageRank scores, color-mapped GO aspects, and a hierarchical display of the GO subgraph. Relationships between GO terms are portrayed with edges and clicking on any node triggers an event handler that populates both the GO information and DEG data to tables.

### Mechanistic Functions for Relationship Mapping

Mechanistically, functions like Parentfinder, Childfinder, SupertermIdentifier, SubtermIdentifier, and AllPathsToRoot map nodes to parent or child GO terms, identify super- and sub-terms, and provide all nodes along the most direct linear route to a given node’s root ontology term. The AllPathsToRoot function provides all nodes along the most direct linear route to a given node’s root ontology term; that is Biological Process (BP), Molecular Function (MF), or Cellular Component (CC). These functions support the NetworkMapper, which utilizes an input node to access specific annotated relationships in the network graph and returns a dictionary with nodes mapped to the specified relationship annotations.

### Utilizing Reference Databases - GPAD, GPI, and UniProt

Combining GPAD and GPI files, a pandas [30] dataframe is generated and used to create a dictionary mapping GO terms to UniProt identifiers [31]. Two distinct lambda functions facilitate conversions between GO term identifiers, UniProt identifiers, and HGNC gene identifiers. Collectively, these functions enable HGNCsymbols_for_go_id to retrieve the HGNC symbols related to a given GO term identifier.

### DEG Association and Output Generation

DEGassociator processes user-supplied file paths and a list of GO terms and outputs the specific DEGs found in the user supplied DEG export files. Matching genes generate a dictionary with GO terms as keys and corresponding dataframe rows. The output_to_csv function concatenates dictionaries into a dataframe, facilitating the generation of a CSV output file.

### Summary and Web Interface

In summary, the main function reads user inputs, generates a list of GO terms through ConcatKeywordHits, processes DEG export files with DEGassociator, and exports result .csv files with output_to_csv. The accompanying server allows users to download a GO network visualization and the GO-mapped DEG CSV file through buttons on the web-based graphical user interface.

### PRRGO User Workflow

Beginning from a keyword query of interest and GSE study for PRRGO analysis, the user would first input the GSE ID into GEO2R and select the treatment and control groups for differential gene expression analysis. The resulting DEG TSV file serves as input to PRRGO. The PRRGO output allows viewing the DEG statistics generated by GEO2R in combination with the GO network image, GO term information, and DEG statistics (Figure 2). The network can be explored using the web interface as well as the output files. While many use cases are possible for the output data, PRRGO lends itself particularly to hypothesis generation, GO-to-DEG mapping, and overlap studies.

**Fig. 2:**
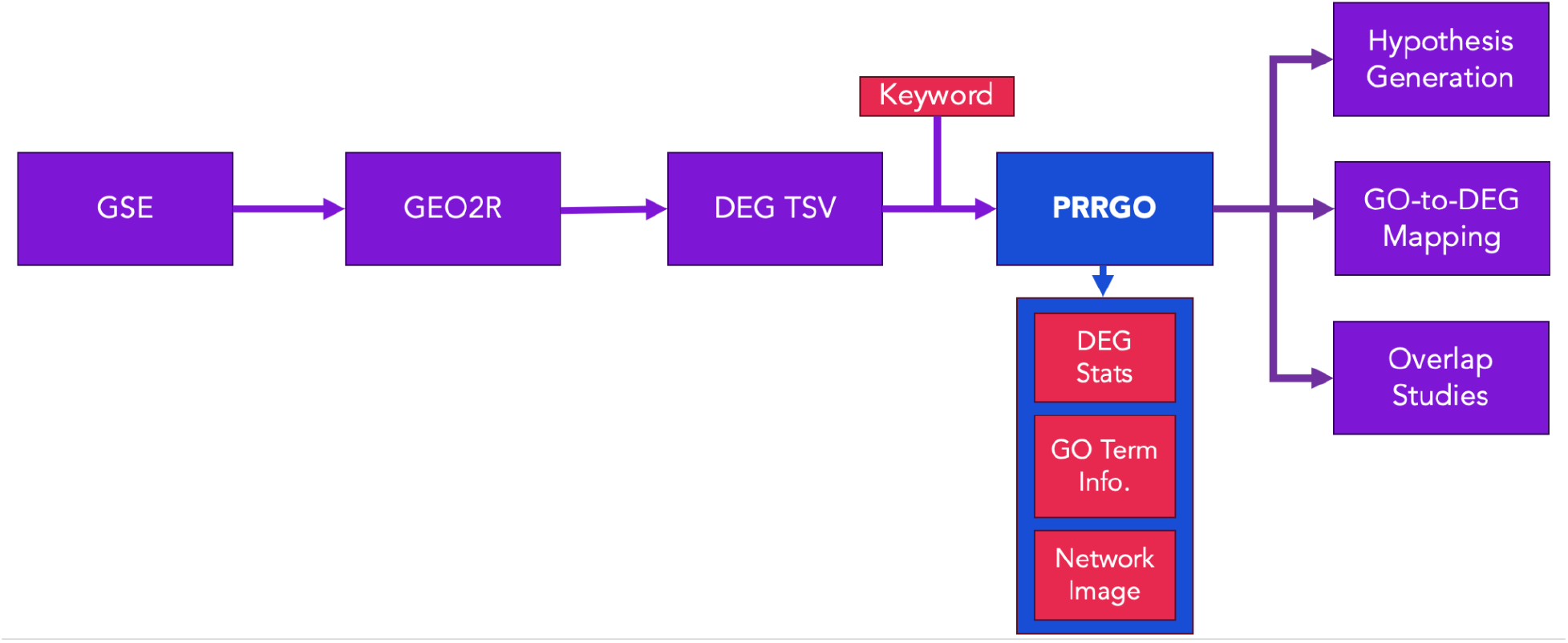
PRRGO User Workflow. The user inputs a GSE ID of interest into GEO2R to generate the DEG TSV. With this TSV as input, PRRGO will output DEG statistics, GO term information (GO domain, description, PageRank), and the GO network image. PRRGO output particularly lends itself to network-based hypothesis generation, GO-to-DEG mapping, and overlap studies.

### Validation Protocol

Candidate GEO Series (GSE) studies containing NCBI-generated raw counts were identified by searching NCBI DataSets with the query “human heat shock ‘rnaseq counts’[Filter]”. Each returned study was iteratively examined for an experimental design wherein gene expression using mRNA sequencing was measured in human cells for at least 2 biological replicates for both control and heat-stress conditions. Nine GSE series met these criteria. Specific samples for inclusion into the validation were defined as representative of either “Control” or “Heat Shock” samples. Detailed information for each GSE is provided in “Sample Inclusion Criteria” (Supplementary file S1). For each of these samples, the GSE identifier was input into the GEO2R web interface. (Alternatively, the “Analyze with GEO2R” option, can be selected from the GEO Accession display from the accession page for a given GSE of interest.) For each GSM identifier meeting inclusion criteria, groups were defined as “Control” or “Heat Shock”. No default options were changed within the options tab and the analyze button was selected. Once finished, the select columns link was selected and the “Check all” box toggled “on” and “set”. Lastly, the “Download full table” link was selected to generate the NCBI-generated DESeq2 analyzed DEG statistics.

For each GSE’s DEG statistics, PRRGO was executed using the search query “Heat shock”. Each GSE examined would yield CSV output files containing GO terms organized by the specific DEGs to which they can be mapped and a representative image of the visualized GO network appearing in PRRGO after entering the indicated query. Lastly, our validation protocol notes considerable parallels between our workflow and the published best practices protocol [32] for analysis of public datasets; steps 1-4 of this published work are effectively streamlined by the use of GEO2R given an existing GSE study identifier within the GEO. Lastly, we provide an R markdown generated .html document to facilitate reproducibility and transparency (Supplementary S4).

## RESULTS

Regarding the study design of the validation, we acknowledge the statistically challenging medium presented by attempting a meta-analysis on inherently stochastic and dispersed expression data [33] of multiple independent studies [34]. The front-end use of GEO2R to standardize the DEG analysis of raw expression data helps mitigate variance due to differences in bioinformatic pipelines used in the original studies. Our justification for use of the canonical heat shock response (HSR) for our validation is multifaceted. HSR is a well-characterized pathway, abundant in supporting literature [35-38]. We reasoned GO term annotations would be accurate, specific, and interpretable. Additionally, the deep evolutionary conservation [38] of HSR means orthogonal experimental validations in virtually any model organism, or representative cell lines thereof, are feasible.

### Validation Summary

In total, 144 samples were collected from nine publicly available datasets, categorized as controls (95) or heat-shocked samples (49), processed with NCBI’s GEO2R website, and then analyzed with PRRGO as described in “GEO2R-to-PRRGO workflow” (Supplementary file S2).

Four samples from study GSE124609 [39] were processed with GEO2R, which reports that 15427 total genes had measurable expression (Table 1). Of these detected genes, 342 protein coding genes were calculated as positively differentially expressed and 552 protein coding genes were calculated as negatively differentially expressed for a total of 894 protein coding genes at statistically significant (p.adj <0.05) levels (Table 1). Of the statistically significant positively differentially expressed genes, 181 exhibited log2FC expression >1 and 227 exhibited log2FC expression <-1 for a total of 408 genes with |log2FC expression|>1 (Table 1). Of the statistically significant differentially expressed genes, GEO2R also reports 970 associated Gene Ontology-Molecular Function terms, 3546 associated Gene Ontology-Biological Process terms and 594 associated Gene Ontology-Cellular Component terms (Table 1). These enriched GO terms contained key HSR indicator GO terms: GO:0031072, GO:0030544, GO:0051879, GO:0140468, GO:0101031.

**Table 1.** Summary statistics performed on GEO2R output of nine human cell line GSE studies involving heat-shock comprise the PRRGO validation.

| GSE Study | Study Information |  |  | Significant DEG GO Terms |  |  |
| --- | --- | --- | --- | --- | --- | --- |
|  | Expt:Control | Detected Genes | DEGs (padj<0.05) | Significant DEG GO:MF terms | Significant DEG GO:BP terms | Significant DEG GO:CC terms |
| GSE 66448 | 2:2 | 17755 | 3140 | 2014 | 6243 | 1093 |
| GSE 164834 | 6:3 | 18042 | 4000 | 2328 | 7381 | 1177 |
| GSE 73471 | 3:2 | 17416 | 1672 | 1483 | 4965 | 855 |
| GSE 123980 | 4:2 | 22115 | 3691 | 2302 | 7202 | 1169 |
| GSE 124510 | 6:47 | 17896 | 7213 | 3085 | 9172 | 1464 |
| GSE 124609 | 4:4 | 15427 | 894 | 970 | 3546 | 594 |
| GSE132447 | 9:3 | 23202 | 2448 | 1793 | 6126 | 918 |
| GSE159802 | 3:6 | 16385 | 2521 | 1715 | 5902 | 952 |
| GSE182960 | 12:26 | 16773 | 5105 | 2598 | 7926 | 1299 |

Nine samples from study GSE123980 [40] were processed with GEO2R, which reports that 22115 total genes had measurable expression (Table 1). Of these detected genes, 1714 protein coding genes were calculated as positively differentially expressed and 1977 protein coding genes were calculated as negatively differentially expressed for a total of 3691 protein coding genes at statistically significant (p.adj <0.05) levels (Table 1). Of the statistically significant positively differentially expressed genes, 52 exhibited log2FC expression >1 and 173 exhibited log2FC expression <-1 for a total of 225 genes with |log2FC expression|>1 (Table 1). Of the statistically significant differentially expressed genes, GEO2R also reports 2302 associated Gene Ontology-Molecular Function terms, 7202 associated Gene Ontology-Biological Process terms and 1169 associated Gene Ontology-Cellular Component terms (Table 1). These enriched GO terms contained key HSR indicator GO terms: GO:0030544, GO:0031072, GO:0051879, GO:0140468, GO:0101031, GO:0072380.

Five samples from study GSE73471 [41] were processed with GEO2R, which reports that 17416 total genes had measurable expression (Table 1). Of these detected genes, 836 protein coding genes were calculated as positively differentially expressed and 836 protein coding genes were calculated as negatively differentially expressed for a total of 1672 protein coding genes at statistically significant (p.adj <0.05) levels (Table 1). Of the statistically significant positively differentially expressed genes, 192 exhibited log2FC expression >1 and 321 exhibited log2FC expression <-1 for a total of 513 genes with |log2FC expression|>1 (Table 1). Of the statistically significant differentially expressed genes, GEO2R also reports 1483 associated Gene Ontology-Molecular Function terms, 4965 associated Gene Ontology-Biological Process terms and 855 associated Gene Ontology-Cellular Component terms (Table 1). These enriched GO terms contained key HSR indicator GO terms: GO:0031072, GO:0030544, GO:0051879, GO:0140468, GO:0101031.

Six samples from study GSE164834 [42] were processed with GEO2R, which reports that 18042 total genes had measurable expression (Table 1). Of these detected genes, 2147 protein coding genes were calculated as positively differentially expressed and 1853 protein coding genes were calculated as negatively differentially expressed for a total of 4000 protein coding genes at statistically significant (p.adj <0.05) levels (Table 1). Of the statistically significant positively differentially expressed genes, 479 exhibited log2FC expression >1 and 39 exhibited log2FC expression <-1 for a total of 518 genes with |log2FC expression|>1 (Table 1). Of the statistically significant differentially expressed genes, GEO2R also reports 2328 associated Gene Ontology-Molecular Function terms, 7381 associated Gene Ontology-Biological Process terms and 1177 associated Gene Ontology-Cellular Component terms (Table 1). These enriched GO terms contained key HSR indicator GO terms: GO:0031072, GO:0030544, GO:0051879, GO:0101031.

Fifty-two samples from study GSE66448 [43] were processed with GEO2R, which reports that 17755 total genes had measurable expression (Table 1). Of these detected genes, 1547 protein coding genes were calculated as positively differentially expressed and 1593 protein coding genes were calculated as negatively differentially expressed for a total of 3140 protein coding genes at statistically significant (p.adj <0.05) levels (Table 1). Of the statistically significant positively differentially expressed genes, 985 exhibited log2FC expression >1 and 772 exhibited log2FC expression <-1 for a total of 1757 genes with |log2FC expression|>1 (Table 1). Of the statistically significant differentially expressed genes, GEO2R also reports 2014 associated Gene Ontology-Molecular Function terms, 6243 associated Gene Ontology-Biological Process terms and 1093 associated Gene Ontology-Cellular Component terms (Table 1). These enriched GO terms contained key HSR indicator GO terms: GO:0031072, GO:0030544, GO:0051879, GO:0140468, GO:0101031.

Eight samples from study GSE124510 [44] were processed with GEO2R, which reports that 17896 total genes had measurable expression (Table 1). Of these detected genes, 3075 protein coding genes were calculated as positively differentially expressed and 4138 protein coding genes were calculated as negatively differentially expressed for a total of 7213 protein coding genes at statistically significant (p.adj <0.05) levels (Table 1). Of the statistically significant positively differentially expressed genes, 1631 exhibited log2FC expression >1 and 2230 exhibited log2FC expression <-1 for a total of 3861 genes with |log2FC expression|>1 (Table 1). Of the statistically significant differentially expressed genes, GEO2R also reports 3085 associated Gene Ontology-Molecular Function terms, 9172 associated Gene Ontology-Biological Process terms and 1464 associated Gene Ontology-Cellular Component terms (Table 1). These enriched GO terms contained key HSR indicator GO terms: GO:0030544, GO:0051879, GO:0031072, GO:0140468, GO:0101031, GO:0072380.

Ten samples from study GSE132447 [45]were processed with GEO2R, which reports that 23202 total genes had measurable expression (Table 1). Of these detected genes, 1527 protein coding genes were calculated as positively differentially expressed and 921 protein coding genes were calculated as negatively differentially expressed for a total of 2448 protein coding genes at statistically significant (p.adj <0.05) levels (Table 1). Of the statistically significant positively differentially expressed genes, 977 exhibited log2FC expression >1 and 273 exhibited log2FC expression <-1 for a total of 1250 genes with |log2FC expression|>1 (Table 1). Of the statistically significant differentially expressed genes, GEO2R also reports 1793 associated Gene Ontology-Molecular Function terms, 6126 associated Gene Ontology-Biological Process terms and 918 associated Gene Ontology-Cellular Component terms (Table 1). These enriched GO terms contained key HSR indicator GO terms: GO:0031072, GO:0030544, GO:0051879, GO:0140468, GO:0101031.

Nine samples from study GSE159802 [46] were processed with GEO2R, which reports that 16385 total genes had measurable expression (Table 1). Of these detected genes, 1181 protein coding genes were calculated as positively differentially expressed and 1340 protein coding genes were calculated as negatively differentially expressed for a total of 2521 protein coding genes at statistically significant (p.adj <0.05) levels (Table 1). Of the statistically significant positively differentially expressed genes, 672 exhibited log2FC expression >1 and 296 exhibited log2FC expression <-1 for a total of 968 genes with |log2FC expression|>1 (Table 1). Of the statistically significant differentially expressed genes, GEO2R also reports 1715 associated Gene Ontology-Molecular Function terms, 5902 associated Gene Ontology-Biological Process terms and 952 associated Gene Ontology-Cellular Component terms (Table 1). These enriched GO terms contained key HSR indicator GO terms: GO:0031072, GO:0030544, GO:0051879, GO:0051008, GO:0101031.

Twenty-four samples from study GSE182960 [47] were processed with GEO2R, which reports that 16773 total genes had measurable expression (Table 1). Of these detected genes, 2640 protein coding genes were calculated as positively differentially expressed and 2465 protein coding genes were calculated as negatively differentially expressed for a total of 5105 protein coding genes at statistically significant (p.adj <0.05) levels (Table 1). Of the statistically significant positively differentially expressed genes, 361 exhibited log2FC expression >1 and 25 exhibited log2FC expression <-1 for a total of 386 genes with |log2FC expression|>1 (Table 1). Of the statistically significant differentially expressed genes, GEO2R also reports 2598 associated Gene Ontology-Molecular Function terms, 7926 associated Gene Ontology-Biological Process terms and 1299 associated Gene Ontology-Cellular Component terms (Table 1). These enriched GO terms contained key HSR indicator GO terms: GO:0030544, GO:0031072, GO:0051879, GO:0051008, GO:0140468, GO:0101031.

## DISCUSSION

Here, we introduce PRRGO, a novel tool designed to simultaneously explore DEG and GO terms. PRRGO is specifically tailored for use in conjunction with NCBI’s GEO2R web application, creating a streamlined pipeline for standardized and reproducible data processing from publicly accessible GEO databases.

PRRGO expeditiously conducts a PageRank-based search, allowing users to specify a keyword or phrase for exploration throughout GO. The search queries GO terms and their descriptions for exact matches, calculating PageRank scores for the resulting GO network (Table 2, Figure 3C). This metric serves as a relative importance indicator for each GO term, enabling comparisons across multiple PRRGO queries or with other tools. However, users are cautioned that this ranking does not incorporate enrichment data and cross-referencing with external enrichment data is recommended for a more comprehensive interpretation.

**Table 2.**
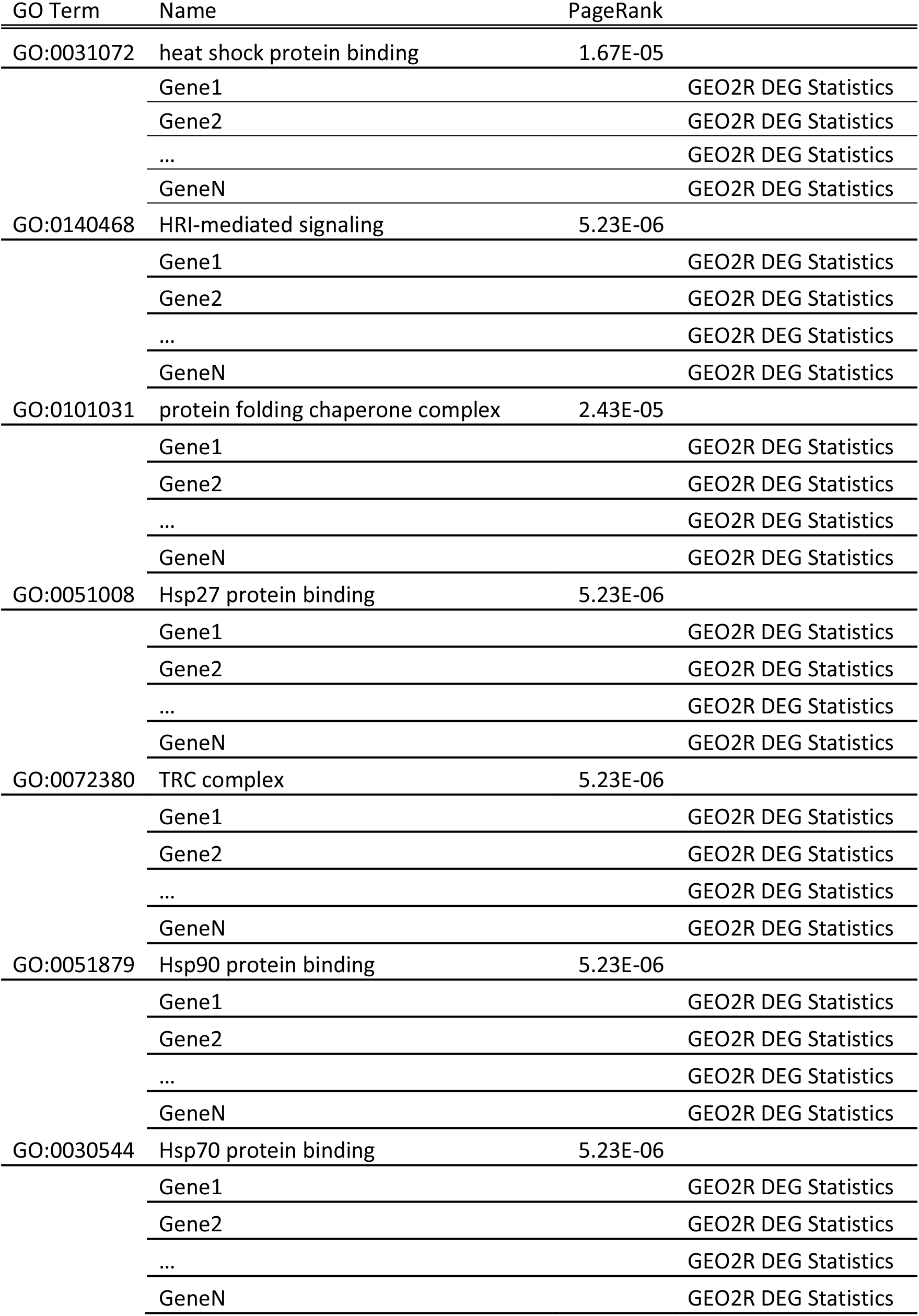
A representative PRRGO output demonstrates that for each keyword-hit GO term, its GO PR score is reported in addition to GEO2R calculated summary DEG statistics for related genes detected.

**FIG. 3:**
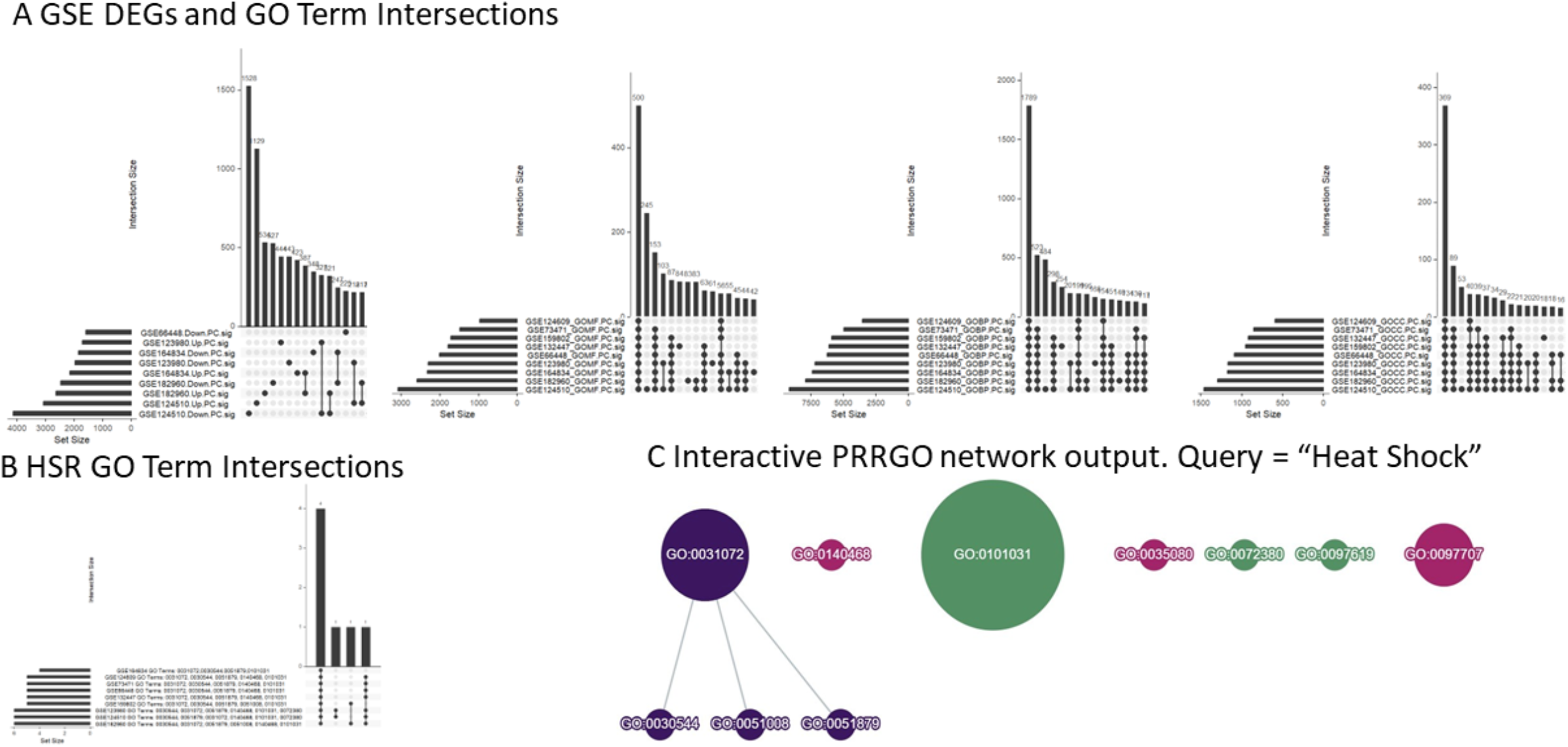
HEAT SHOCK RESPONSE GO TERMS IDENTIFIED IN PUBLIC GEO DATASETS VALIDATE PRRGO FUNCTIONALITY. **A.** UpSet plots exhibit higher concordance intersections among enriched GO terms than in DEGs within the selected GSE studies. **B**. Upset plots demonstrate HSR specific GO Terms enriched in selected GSE studies are virtually unanimously concordant. **C**. A representative PRRGO network output generated by the query “heat shock” communicates a PR score (node size), GO source (purple:MF, magenta:BP, green:CC), and hierarchical relationships (edges).

An essential function of PRRGO is its ability to display DEG statistics from GEO2R for all genes annotated to a specific GO term (Table 2). This functionality addresses the common scenario where researchers possess a list of DEGs for enrichment or pathway analyses. PRRGO complements existing tools [20, 23-27] by facilitating hypothesis generation in the reverse direction—starting from a known biological process, molecular function, or cellular component of interest within the Gene Ontology. The graphical representation of the GO network graph (Figure 3C), along with the DEG table (Table 2), allows users to efficiently explore GO terms related to a keyword of interest. It is important to note that the DEG statistics table is displayed for all genes with GO annotations, regardless of DEG significance or GO enrichment status.

PRRGO enhances exploration efficiency by combining a hierarchical GO network graph representation with a detailed DEG table. The DEG statistics table is displayed for all genes with GO annotations, providing a comprehensive overview regardless of DEG significance or GO enrichment status. Additionally, PRRGO facilitates the export of publication-ready network representations of GO term hits (Figure 3C), generating PNG images and summary CSV files. The CSV file is particularly valuable for curating inputs to downstream analysis tools, leveraging the wealth of DEG lists and associated GO term annotations.

PRRGO’s internal architecture leverages data structures from public data, enabling the use of any public GSE study interpretable by GEO2R for validation. Public data sourced from peer-reviewed studies ensures adherence to a minimum standard of sample quality and experimental rigor, instilling confidence in the tool’s reliability and relevance.

### Intended Use and Scope

PRRGO serves as a powerful exploratory tool designed to parse and connect GO and DEG data (Figure 3B, Figure 4). While it does not replace traditional enrichment analysis tools, PRRGO excels in providing a user-friendly means of descriptive and exploratory analysis. Derived mainly from external sources, the tool visualizes data in a human-readable format, facilitating rapid network-based hypothesis generation from richly annotated gene expression databases. PRRGO’s versatility is highlighted in its ability to explore the overlap of multiple GO terms and generate diverse plots for traditional DEG analysis, incorporating PageRank data for enhanced comparison across studies and GO terms (Figures 4, Figure 5).

**Fig. 4:**
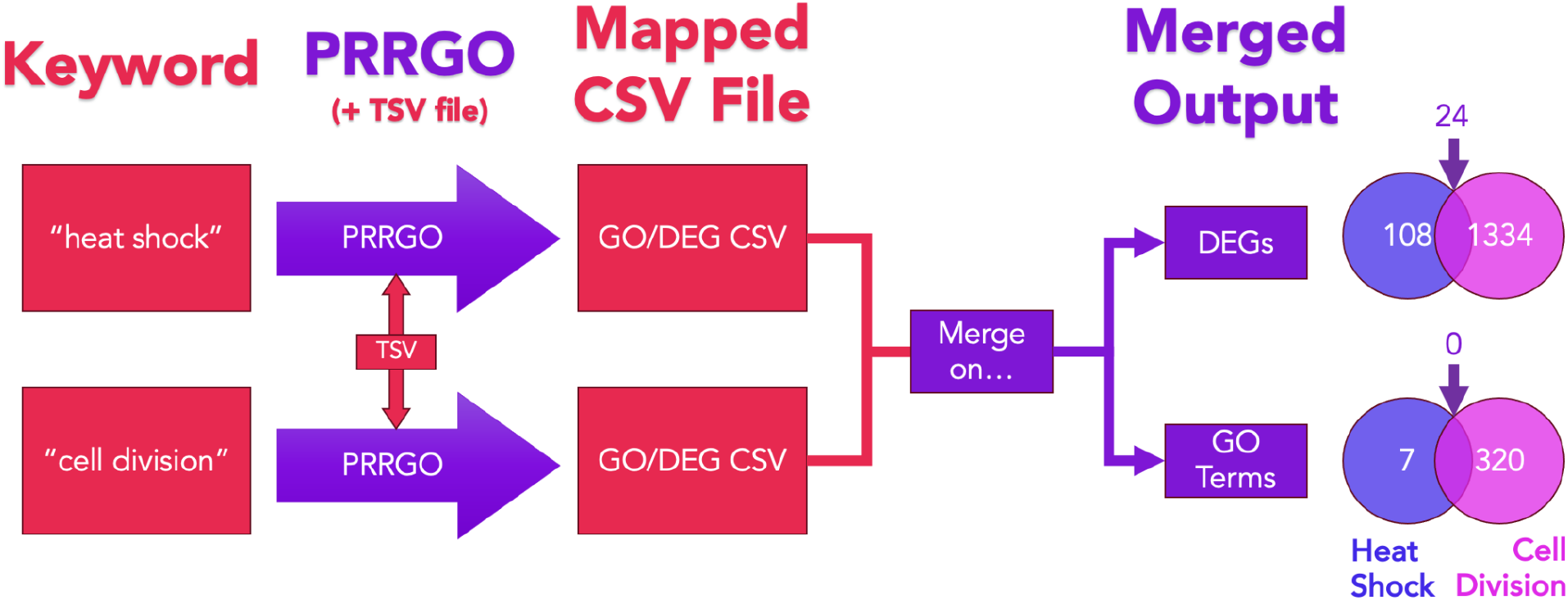
Sample GO Term/DEG Intersection Use Case. After querying PRRGO with the keywords “heat shock” and “cell division”, the resulting GO-mapped DEG CSV files were merged. Then, the overlap between the GO terms and DEGs was found and visualized with Venn diagrams.

**Fig. 5:**
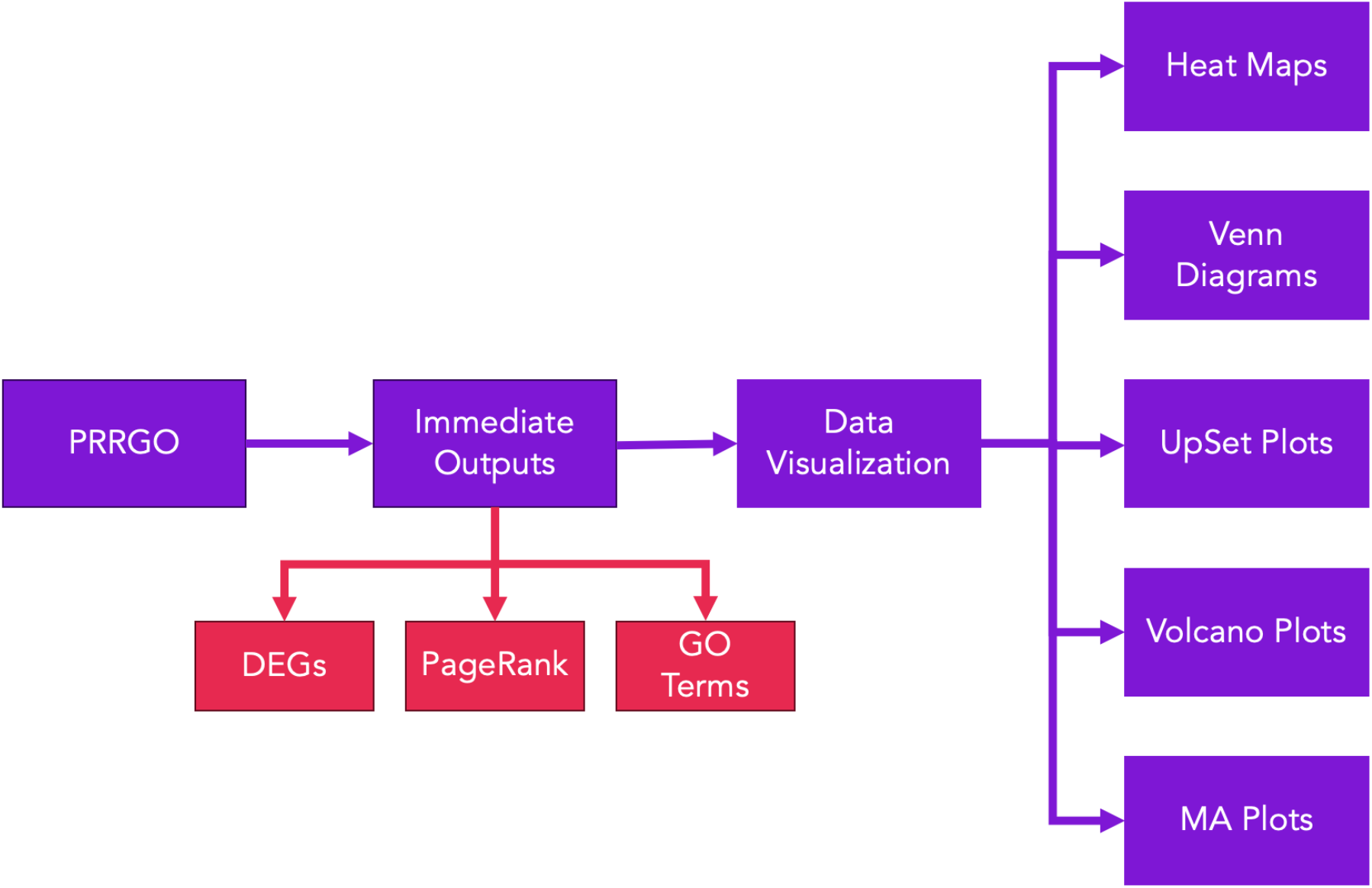
Summary of Potential Figures. In conjunction with other data visualization tools, PRRGO’s output may be used to generate a variety of figures: heat maps, Venn diagrams, UpSet plots, volcano plots, and MA plots.

### Prospective Applications

We envision several potential directions for PRRGO’s development. The simplest yet impactful addition could be expanding the accepted input DEG data to accommodate files from various popular DEG analysis tools, such as DESeq2 [3], limma [5], and edgeR [4]. The flexibility of these tools, while beneficial to users, poses an integration challenge due to differences in formatting. Should this functionality be added, PRRGO’s mapping utility would scale across different workflows seamlessly.

Additionally, incorporating additional quantitative visualizations, especially involving both DEG and GO enrichment data, would enhance PRRGO’s utility. This improvement would enable researchers to generate publication-ready figures, including heat maps, with ease. Customizable outputs could further empower users to selectively choose essential information for subsequent analysis, catering to diverse research needs.

Furthermore, integration with tools and databases like String [48] and Cytoscape [21] would extend PRRGO’s potential utility. Lastly, adding the capability to parse GO data from species other than Homo sapiens would be particularly valuable, considering that GO contains annotations for various species, even if GEO2R is limited to human data.

## Conclusion

PRRGO functions as an efficient GO-DEG exploratory web application that can be used by biologists with internet access for GEO2R and familiarity with the command line. Its ability to integrate GO and DEG data and display in both the easily navigable, interactive web format and the output CSV file greatly expedites DEG data exploration. Its novel use of the PageRank algorithm to show the relative seniority of GO terms provides a simple method for prioritizing more senior GO terms of interest.

## Supporting information

S1_Sample_Inclusion_Criteria

S2_GEO2R-PRRGO_Complete_Workflow

S3_Validation_Inputs-Outputs

S4_DEGMetaAnalysis

## Data Availability

PRRGO is publicly available at https://github.com/ndsa23/PRRGO under a GNU Lesser General Public License v2.1. Detailed installation instructions can be found in the README.md in the Github repository. PRRGO is platform-independent and can run on Windows, MacOS, and Linux on all major browsers.

## Supplementary Material

The sample inclusion criteria for all heat shock studies are available in supplementary material S1.

The complete GEO2R to PRRGO workflow is available in supplementary material S2.

The PRRGO heat shock studies output is available in supplementary material S3.

The heat shock studies analysis R markdown HTML file is available in supplementary material S4.

## Acknowledgements

We thank the members of the Nikolaidis group, especially Andrew Reinschmidt and Yonny Chavez, for their feedback.

## References

1. Ashburner M, Ball CA, Blake JA, Botstein D, Butler H, Cherry JM, Davis AP, Dolinski K, Dwight SS, Eppig JT et al: Gene Ontology: tool for the unification of biology. Nature Genetics 2000, 25(1):25–29.

2. Barrett T, Wilhite SE, Ledoux P, Evangelista C, Kim IF, Tomashevsky M, Marshall KA, Phillippy KH, Sherman PM, Holko M et al: NCBI GEO: archive for functional genomics data sets—update. Nucleic Acids Research 2012, 41(D1):D991–D995.

3. Love MI, Huber W, Anders S: Moderated estimation of fold change and dispersion for RNA-seq data with DESeq2. Genome Biology 2014, 15(12):550.

4. Robinson MD, McCarthy DJ, Smyth GK: edgeR: a Bioconductor package for differential expression analysis of digital gene expression data. Bioinformatics 2009, 26(1):139–140.

5. Ritchie ME, Phipson B, Wu D, Hu Y, Law CW, Shi W, Smyth GK: limma powers differential expression analyses for RNA-sequencing and microarray studies. Nucleic Acids Research 2015, 43(7):e47–e47.

6. Law CW, Chen Y, Shi W, Smyth GK: voom: precision weights unlock linear model analysis tools for RNA-seq read counts. Genome Biology 2014, 15(2):R29.

7. Seyednasrollah F, Laiho A, Elo LL: Comparison of software packages for detecting differential expression in RNA-seq studies. Briefings in Bioinformatics 2015, 16(1):59–70.

8. The Gene Ontology C, Aleksander SA, Balhoff J, Carbon S, Cherry JM, Drabkin HJ, Ebert D, Feuermann M, Gaudet P, Harris NL et al: The Gene Ontology knowledgebase in 2023. Genetics 2023, 224(1):iyad031.

9. Brin S, Page L: The anatomy of a large-scale hypertextual Web search engine. Computer Networks and ISDN Systems 1998, 30(1):107–117.

10. Page L, Brin S, Motwani R, Winograd T: The PageRank citation ranking: Bringing order to the Web. In. Stanford Digital Library Working Paper SIDL-WP-1999-0120: Stanford University; 1999.

11. Kimmel C, Visweswaran S: An Algorithm for Network-Based Gene Prioritization That Encodes Knowledge Both in Nodes and in Links. PLOS ONE 2013, 8(11):e79564.

12. Zeng X, Zhao J, Wu X, Shi H, Liu W, Cui B, Yang L, Ding X, Song P: PageRank analysis reveals topologically expressed genes correspond to psoriasis and their functions are associated with apoptosis resistance. Mol Med Rep 2016, 13(5):3969–3976.

13. Jiang B, Kloster K, Gleich DF, Gribskov M: AptRank: an adaptive PageRank model for protein function prediction on bi-relational graphs. Bioinformatics 2017, 33(12):1829–1836.

14. Morrison JL, Breitling R, Higham DJ, Gilbert DR: GeneRank: Using search engine technology for the analysis of microarray experiments. BMC Bioinformatics 2005, 6(1):233.

15. Li J, Zhao PX: Mining Functional Modules in Heterogeneous Biological Networks Using Multiplex PageRank Approach. Frontiers in Plant Science 2016, 7.

16. Zhao Q, Zhang Y, Zhang X, Sun Y, Lin Z: Mining of gene modules and identification of key genes in head and neck squamous cell carcinoma based on gene co-expression network analysis. Medicine 2020, 99(49):e22655.

17. Lamurias A, Ruas P, Couto FM: PPR-SSM: personalized PageRank and semantic similarity measures for entity linking. BMC Bioinformatics 2019, 20(1):534.

18. Ding H, Yang Y, Xue Y, Seninge L, Gong H, Safavi R, Califano A, Stuart JM: Prioritizing transcriptional factors in gene regulatory networks with PageRank. iScience 2021, 24(1).

19. Kaalia R, Ghosh I: Semantics based approach for analyzing disease-target associations. Journal of Biomedical Informatics 2016, 62:125–135.

20. Bindea G, Mlecnik B, Hackl H, Charoentong P, Tosolini M, Kirilovsky A, Fridman W-H, Pagès F, Trajanoski Z, Galon J: ClueGO: a Cytoscape plug-in to decipher functionally grouped gene ontology and pathway annotation networks. Bioinformatics 2009, 25(8):1091–1093.

21. Shannon P, Markiel A, Ozier O, Baliga NS, Wang JT, Ramage D, Amin N, Schwikowski B, Ideker T: Cytoscape: a software environment for integrated models of biomolecular interaction networks. Genome Res 2003, 13(11):2498–2504.

22. Kanehisa M, Goto S: KEGG: Kyoto Encyclopedia of Genes and Genomes. Nucleic Acids Research 2000, 28(1):27–30.

23. Pomaznoy M, Ha B, Peters B: GOnet: a tool for interactive Gene Ontology analysis. BMC Bioinformatics 2018, 19(1):470.

24. Wei Q, Khan IK, Ding Z, Yerneni S, Kihara D: NaviGO: interactive tool for visualization and functional similarity and coherence analysis with gene ontology. BMC Bioinformatics 2017, 18(1):177.

25. Ye J, Zhang Y, Cui H, Liu J, Wu Y, Cheng Y, Xu H, Huang X, Li S, Zhou A et al: WEGO 2.0: a web tool for analyzing and plotting GO annotations, 2018 update. Nucleic Acids Research 2018, 46(W1):W71–W75.

26. Zhu J, Zhao Q, Katsevich E, Sabatti C: Exploratory Gene Ontology Analysis with Interactive Visualization. Scientific Reports 2019, 9(1):7793.

27. Carbon S, Ireland A, Mungall CJ, Shu S, Marshall B, Lewis S, Hub tA, Group tWPW: AmiGO: online access to ontology and annotation data. Bioinformatics 2008, 25(2):288–289.

28. Hagberg A, Swart PJ, Schult DA: Exploring network structure, dynamics, and function using NetworkX. In: 2008-01-01 2008; United States. Research Org.: Los Alamos National Laboratory (LANL), Los Alamos, NM (United States); Sponsor Org.: USDOE.

29. Franz M, Lopes CT, Huck G, Dong Y, Sumer O, Bader GD: Cytoscape.js: a graph theory library for visualisation and analysis. Bioinformatics 2016, 32(2):309–311.

30. McKinney W: Data Structures for Statistical Computing in Python. In: Proceedings of the 9th Python in Science Conference: 2010. 56–61.

31. The UniProt C: UniProt: the Universal Protein Knowledgebase in 2023. Nucleic Acids Research 2023, 51(D1):D523–D531.

32. Sanchis P, Lavignolle R, Abbate M, Lage-Vickers S, Vazquez E, Cotignola J, Bizzotto J, Gueron G: Analysis workflow of publicly available RNA-sequencing datasets. STAR Protocols 2021, 2(2):100478.

33. Raser JM, O’Shea EK: Noise in gene expression: origins, consequences, and control. Science 2005, 309(5743):2010–2013.

34. Rau A, Marot G, Jaffrézic F: Differential meta-analysis of RNA-seq data from multiple studies. BMC Bioinformatics 2014, 15(1):91.

35. Morimoto RI: Regulation of the heat shock transcriptional response: cross talk between a family of heat shock factors, molecular chaperones, and negative regulators. Genes Dev 1998, 12(24):3788–3796.

36. Shamovsky I, Ivannikov M, Kandel ES, Gershon D, Nudler E: RNA-mediated response to heat shock in mammalian cells. Nature 2006, 440(7083):556–560.

37. Richter K, Haslbeck M, Buchner J: The Heat Shock Response: Life on the Verge of Death. Molecular Cell 2010, 40(2):253–266.

38. Mahat Dig B, Salamanca HH, Duarte Fabiana M, Danko Charles G, Lis John T: Mammalian Heat Shock Response and Mechanisms Underlying Its Genome-wide Transcriptional Regulation. Molecular Cell 2016, 62(1):63–78.

39. Sabath N, Levy-Adam F, Younis A, Rozales K, Meller A, Hadar S, Soueid-Baumgarten S, Shalgi R: Cellular proteostasis decline in human senescence. Proc Natl Acad Sci U S A 2020, 117(50):31902–31913.

40. Gressel S, Schwalb B, Cramer P: The pause-initiation limit restricts transcription activation in human cells. Nat Commun 2019, 10(1):3603.

41. Hussong M, Kaehler C, Kerick M, Grimm C, Franz A, Timmermann B, Welzel F, Isensee J, Hucho T, Krobitsch S et al: The bromodomain protein BRD4 regulates splicing during heat shock. Nucleic Acids Res 2017, 45(1):382–394.

42. Gibellini L, Borella R, De Gaetano A, Zanini G, Tartaro DL, Carnevale G, Beretti F, Losi L, De Biasi S, Nasi M et al: Evidence for mitochondrial Lonp1 expression in the nucleus. Sci Rep 2022, 12(1):10877.

43. Niskanen EA, Malinen M, Sutinen P, Toropainen S, Paakinaho V, Vihervaara A, Joutsen J, Kaikkonen MU, Sistonen L, Palvimo JJ: Global SUMOylation on active chromatin is an acute heat stress response restricting transcription. Genome Biol 2015, 16(1):153.

44. Huang HH, Ferguson ID, Thornton AM, Bastola P, Lam C, Lin YT, Choudhry P, Mariano MC, Marcoulis MD, Teo CF et al: Proteasome inhibitor-induced modulation reveals the spliceosome as a specific therapeutic vulnerability in multiple myeloma. Nat Commun 2020, 11(1):1931.

45. Bhattacharya A, Jha V, Singhal K, Fatima M, Singh D, Chaturvedi G, Dholakia D, Kutum R, Pandey R, Bakken TE et al: Multiple Alu Exonization in 3’UTR of a Primate-Specific Isoform of CYP20A1 Creates a Potential miRNA Sponge. Genome Biol Evol 2021, 13(1).

46. Vydra N, Janus P, Kus P, Stokowy T, Mrowiec K, Toma-Jonik A, Krzywon A, Cortez AJ, Wojtas B, Gielniewski B et al: Heat shock factor 1 (HSF1) cooperates with estrogen receptor α (ERα) in the regulation of estrogen action in breast cancer cells. Elife 2021, 10.

47. Singh AK, Chen Q, Nguyen C, Meerzaman D, Singer DS: Cohesin regulates alternative splicing. Sci Adv 2023, 9(9):eade3876.

48. Szklarczyk D, Franceschini A, Wyder S, Forslund K, Heller D, Huerta-Cepas J, Simonovic M, Roth A, Santos A, Tsafou KP et al: STRING v10: protein-protein interaction networks, integrated over the tree of life. Nucleic Acids Res 2015, 43(Database issue):D447–452.

