## Supplementary material for "PRRGO: A Tool for Visualizing and Mapping Globally Expressed Genes in Public Gene Expression Omnibus RNA-Sequencing Studies to PageRank-scored Gene Ontology Terms": S1_Sample_Inclusion_Criteria

In GSE124609, RNA-seq analysis was performed on 2 technical replicates of primary human fibroblast cells with 2 heat shock groups (HS at 44 °C for 2 h) and 2 control groups both with young and senescent cells. About 550 genes were differentially expressed upon HS or between young and senescent cells based on differential expression analysis. The following gene ontology (GO) terms were found to be enriched in the proteostasis decline cluster of senescent HS: GO:0006986 (response to unfolded protein), GO:0051082 (unfolded protein binding), GO:0042026 (protein refolding), GO:0051087 (chaperone binding), GO:1900034 (regulation of cellular response to heat), GO:0006457 (protein folding), GO:0031396 (regulation of protein ubiquitination), GO:0031072 (heat shock protein binding), and GO:0009408 (response to heat). The senescence-enhanced HS repression cluster was found to have the following GO terms enriched: GO:0005886 (plasma membrane), GO:0005887 (integral component of plasma membrane), GO:0007229 (integrin-mediated signaling pathway), GO:0016021 (integral component of membrane), GO:0043547 (positive regulation of GTPase activity), GO:0007165 (signal transduction).
For GEO2R analysis, both young and senescent heat shock groups were grouped together as the heat shock group while the young and senescent cells that were not subjected to heat shock were chosen as controls.

| Designation | Availability | Sample ID | Sample description |
|---|---|---|---|
| Control | Yes | GSM3537582 | WI38_young_Control_R1 |
| Control | Yes | GSM3537583 | WI38_young_Control_R2 |
| Heat Shock | Yes | GSM3537584 | WI38_young_HS_R1 |
| Heat Shock | Yes | GSM3537585 | WI38_young_HS_R2 |
| Control | Yes | GSM3537586 | WI38_senescent_Control_R3 |
| Control | Yes | GSM3537587 | WI38_senescent_Control_R4 |
| Heat Shock | Yes | GSM3537588 | WI38_senescent_HS_R3 |
| Heat Shock | Yes | GSM3537589 | WI38_senescent_HS_R4 |

In GSE123980, RNA-seq analysis was performed on 2 biological replicates of human K562 erythroleukemia cells. Two heat shock groups included cells heated to 42 °C for a time period of 15 min and 30 min. The one control group was maintained at 37 °C. 899 genes were found to be significantly upregulated while 2614 genes were found to be significantly downregulated based on a combination of RNA-seq, TT-seq, and mNET-seq. The study ran a GO

overrepresentation analysis on an aggregate of the RNA-seq, TT-seq and mNET-seq data. The following upregulated GO terms were found to be significantly overrepresented (in supplemental information): Heat shock protein binding, cellular response to heat, Hsp70 protein binding, protein refolding, chaperone binding, response to unfolded protein, regulation of cellular response to heat, unfolded protein binding. The following downregulated GO terms were found to be significantly overrepresented (in supplemental information): cell division; mRNA export from nucleus; mRNA processing; mRNA splicing, via spliceosome; rRNA processing; transcription, DNA-templated; poly(A) RNA binding; and nucleoplasm.
For GEO2R analysis, the two heat shock groups were grouped into one heat shock group and compared to the control group. Only RNA-seq results were included in the GEO2R analysis to ensure that the DEGs identified are not confounded with the type of sequencing.

| Designation | Availability | Sample ID | Sample description |
|---|---|---|---|
| TTSeq | excluded | GSM3518105 | control |
| TTSeq | excluded | GSM3518106 | control |
| TTSeq | excluded | GSM3518107 | HeatShock |
| TTSeq | excluded | GSM3518108 | HeatShock |
| TTSeq | excluded | GSM3518109 | HeatShock |
| TTSeq | excluded | GSM3518110 | HeatShock |
| Control | Yes | GSM3518111 | K562_control_R1 |
| Control | Yes | GSM3518112 | K562_control_R2 |
| Heat Shock | Yes | GSM3518113 | K562_HeatShock_R1 |
| Heat Shock | Yes | GSM3518114 | K562_HeatShock_R2 |
| Heat Shock | Yes | GSM3518115 | K562_HeatShock_R3 |
| Heat Shock | Yes | GSM3518116 | K562_HeatShock_R4 |

In GSE66448, GRO-seq analysis was performed on 2 biological replicates of K562 cells with one heat shock group (30 min at 43 °C) and one control group. GO terms were found to be significantly enriched in several gene groups:

- HS-induced: response to unfolded protein, protein folding, programmed cell death, apoptosis, response to heat
- HS-repressed: gene expression, metabolic process, biosynthetic process, transcription, translation, translational elongation, ncRNA metabolic process

For GEO2R analysis, the one heat shock group was included as the heat shock group and the control group was not heat shocked.

| Designation | Availability | Sample ID | Sample description |
|---|---|---|---|
| Control | Yes | GSM1622612 | K562-GROseq-C-rep1 |
| Control | Yes | GSM1622613 | K562-GROseq-C-rep2 |
| Heat Shock | Yes | GSM1622614 | K562-GROseq-HS-rep1 |
| Heat Shock | Yes | GSM1622615 | K562-GROseq-Hs-rep2 |

In GSE73471, RNA-seq was performed on WI38 lung fibroblast cells of heat shock (HS) groups (42 °C for 4 h), control groups, and RNAi targeting BRD4 (gene involved in transcriptional regulation and mRNA splicing of heat shock). For GEO2R analysis, the two non-HS groups (one wild-type and the other RNAi BRD4) were used as controls. The non-RNAi HS group (3 biological replicates) was used as the comparison, so that it could be observed whether canonical heat shock proteins were expressed.

| Designation | Availability | Sample ID | Sample description |
|---|---|---|---|
| Control | Yes | GSM1895365 | Ctrl 1 |
| Control | Yes | GSM1895366 | si BRD4 1 |
| Heat Shock | Yes | GSM1895367 | HS 1 |

| | | | |
|---|---|---|---|
| NA | Yes | GSM1895368 | si BRD4 & HS 1 |
| Heat Shock | Yes | GSM1895369 | HS 2 |
| NA | Yes | GSM1895370 | si BRD4 & HS 2 |
| Heat Shock | Yes | GSM1895371 | HS 3 |
| NA | Yes | GSM1895372 | si BRD4 & HS 3 |

In GSE165367, TT-seq analysis was performed on three biological replicates of heat shock (43 °C for 2 h) and non-HS in MRC5VA cells. Differential gene expression analysis yielded 2,324 downregulated genes and 1,622 upregulated genes.
For GEO2R analysis, the three biological replicates that were heat shocked were treated as the heat shock group and the three non-HS biological replicates were treated as the control.
Control: GSM5032228, GSM5032229, GSM5032230
HS: GSM5032231, GSM5032232, GSM5032233

InGSE124510RNA-seq analysis was performed on multiple myeloma or AMO cells treated withE7107, Carfilzomib,DMSO, melphalan, or bortezomib, none of which have documented effects on the heat shockresponse.Additionally, 3 samples were heat shocked and immediately processed for mRNA-sequencingand 3 samples were heat shocked and allowed to recover at control conditions for 4 hours beforebeing processed for mRNA-seq.These samples were collectively designated heat shocked and allothers were designated controls. 18 samples were WT or mutants for the SRSF1 gene

| Designation | Availability | Sample ID | Sample description |
|---|---|---|---|
| control | yes | GSM3535537 | MM.1S30nMcarfilzomib0hmRNA |
| control | yes | GSM3535538 | MM.1S30nMcarfilzomib8hmRNA |
| control | yes | GSM3535539 | MM.1S30nMcarfilzomib16hmRNA |
| control | yes | GSM3535540 | MM.1S30nMcarfilzomib24hmRNA |
| control | yes | GSM3535541 | MM.1SDMSOmRNAreplicate3 |
| control | yes | GSM3535542 | MM.1SDMSOmRNAreplicate4 |

| | | | |
|---|---|---|---|
| control | yes | GSM3535543 | MM.1SDMSOmRNAreplicate8 |
| control | yes | GSM3535544 | MM.1S18nMcarfilzomibmRNAreplicate1 |
| control | yes | GSM3535545 | MM.1S18nMcarfilzomibmRNAreplicate4 |
| control | yes | GSM3535546 | MM.1S18nMcarfilzomibmRNAreplicate3 |
| control | yes | GSM3535547 | MM.1S10uMmelphalanmRNAreplicate1 |
| control | yes | GSM3535548 | MM.1S10uMmelphalanmRNAreplicate2 |
| control | yes | GSM3535549 | MM.1S10uMmelphalanmRNAreplicate10 |
| control | yes | GSM3535550 | MM.1S10nME7107mRNAreplicate1 |
| control | yes | GSM3535551 | MM.1S10nME7107mRNAreplicate3 |
| control | yes | GSM3535552 | MM.1S10nME7107mRNAreplicate4 |
| control | yes | GSM3535553 | AMO-110nME7107mRNAreplicate1 |
| control | yes | GSM3535554 | AMO-110nME7107mRNAreplicate3 |
| control | yes | GSM3535555 | AMO-110nME7107mRNAreplicate4 |
| control | yes | GSM3535556 | AMO-1DMSOmRNAreplicate4 |
| control | yes | GSM3535557 | AMO-1DMSOmRNAreplicate8 |
| control | yes | GSM3535558 | AMO-1DMSOmRNAreplicate7 |
| control | yes | GSM3535559 | AMO-115nMcarfilzomibmRNAreplicate4 |
| control | yes | GSM3535560 | AMO-115nMcarfilzomibmRNAreplicate8 |
| control | yes | GSM3535561 | AMO-115nMcarfilzomibmRNAreplicate7 |
| control | yes | GSM3535562 | AMO-1SRSF1WTDMSOmRNAreplicate4 |
| control | yes | GSM3535563 | AMO-1SRSF1WTDMSOmRNAreplicate2 |
| control | yes | GSM3535564 | AMO-1SRSF1WTDMSOmRNAreplicate3 |
| control | yes | GSM3535565 | AMO-1SRSF1mSDDMSOmRNAreplicate4 |

| | | | |
|---|---|---|---|
| control | yes | GSM3535566 | AMO-1SRSF1mSDDMSOmRNAreplicate2 |
| control | yes | GSM3535567 | AMO-1SRSF1mSDDMSOmRNAreplicate3 |
| control | yes | GSM3535568 | AMO-1SRSF1mSADMSOmRNAreplicate4 |
| control | yes | GSM3535569 | AMO-1SRSF1mSADMSOmRNAreplicate2 |
| control | yes | GSM3535570 | AMO-1SRSF1mSADMSOmRNAreplicate3 |
| control | yes | GSM3535571 | AMO-1SRSF1WT15nMcarfilzomibmRNA |
| control | yes | GSM3535572 | AMO-1SRSF1WT15nMcarfilzomibmRNA |
| control | yes | GSM3535573 | AMO-1SRSF1WT15nMcarfilzomibmRNA |
| control | yes | GSM3535574 | AMO-1SRSF1mSD15nMcarfilzomibmRNAreplicate |
| control | yes | GSM3535575 | AMO-1SRSF1mSD15nMcarfilzomibmRNAreplicate |
| control | yes | GSM3535576 | AMO-1SRSF1mSD15nMcarfilzomibmRNAreplicate |
| control | yes | GSM3535577 | AMO-1SRSF1mSA15nMcarfilzomibmRNAreplicate |
| control | yes | GSM3535578 | AMO-1SRSF1mSA15nMcarfilzomibmRNAreplicate |
| control | yes | GSM3535579 | AMO-1SRSF1mSA15nMcarfilzomibmRNAreplicate |
| heatshock | yes | GSM4231704 | MM.1S |
| heatshock | yes | GSM4231705 | MM.1S |
| heatshock | yes | GSM4231706 | MM.1S |
| heatshock | yes | GSM4231707 | MM.1S |
| heatshock | yes | GSM4231708 | MM.1S |
| heatshock | yes | GSM4231709 | MM.1S |

| | | | |
|---|---|---|---|
| control | yes | GSM4231710 | MM.1S |
| control | yes | GSM4231711 | MM.1S |
| control | yes | GSM4231712 | MM.1S |

In GSE132447, RNA-seq analysis was performed on 3 biological replicates of primary human neurons following control, or various heat-shock (HS) and recovery conditions (45C for 30 minutes and recovery at 37C for 1 hour, 3 hours, or 24 hours). Several thousand genes (~3K) were found differentially expressed in HS conditions, of which 380 were used as input for a gene ontology (GO) analysis. The top five GO processes mapped to hemostasis (28 genes), axon guidance (25 genes), neutrophil degranulation (23 genes), platelet signaling, activation and aggregation (18 genes), and ECM organization (18 genes). Additional GO findings included mRNA processing and mitochondria translation, metabolism, amino acid and nucleotide synthesis, and antigen presentation.

| Designation | Availability | Sample ID | Sample description |
|---|---|---|---|
| Control | Yes | GSM3864912 | RNA seq_Untreated_Replicate_1 |
| Control | Yes | GSM3864913 | RNA seq_Untreated_Replicate_2 |
| Control | Yes | GSM3864914 | RNA seq_Untreated_Replicate_3 |
| Heat-Shock | Yes | GSM3864915 | RNA seq_Heat_shock_1 hr recovery_Replicate_1 |
| Heat-Shock | No | GSM3864916 | RNA seq_Heat_shock_1 hr recovery_Replicate_2 |
| Heat-Shock | No | GSM3864917 | RNA seq_Heat_shock_1 hr recovery_Replicate_3 |
| Heat-Shock | Yes | GSM3864918 | RNA seq_Heat_shock_3 hr recovery_Replicate_1 |
| Heat-Shock | Yes | GSM3864919 | RNA seq_Heat_shock_3 hr recovery_Replicate_2 |
| Heat-Shock | Yes | GSM3864920 | RNA seq_Heat_shock_3 hr recovery_Replicate_3 |

| | | | |
|---|---|---|---|
| Heat-Shock | Yes | GSM3864921 | RNA seq_Heat_shock_24 hr recovery_Replicate_1 |
| Heat-Shock | Yes | GSM3864922 | RNA seq_Heat_shock_24 hr recovery_Replicate_2 |
| Heat-Shock | Yes | GSM3864923 | RNA seq_Heat_shock_24 hr recovery_Replicate_3 |
| exclude | Yes | GSM3864924 | RNA seq_Tat_no recovery_Replicate_1 |
| exclude | Yes | GSM3864925 | RNA seq_Tat_no recovery_Replicate_2 |
| exclude | Yes | GSM3864926 | RNA seq_Tat_no recovery_Replicate_3 |
| exclude | Yes | GSM3864927 | RNA seq_Tat_6 hr recovery_Replicate_1 |
| exclude | Yes | GSM3864928 | RNA seq_Tat_6 hr recovery_Replicate_2 |
| exclude | Yes | GSM3864929 | RNA seq_Tat_6 hr recovery_Replicate_3 |

19 Samples from GSE159802 were subject to various permutations of estrogen treatments, heat-shock and recovery conditions, or control conditions (37C). Three biological replicates were heat shocked where each differed only in whether they were also treated with a shRNA knock-down control, a CRISPR/Cas9 control, or no other treatment. Six biological replicates we classify as HS control samples can be further categorized as either a shRNA knock-down control, a CRISPR/Cas9 control, estrogen treatment, or estrogen treatment control. Gene set enrichment analysis (GSEA) yielded the following 12 GO terms related to Hormone signaling and metabolism: response to estrogen, response to estradiol, response to hormone, cellular response to hormone stimulus, cellular response to estradiol stimulus, regulation of peptide hormone secretion, positive regulation of hormone secretion, hormone metabolic process, peptide hormone secretion peptide hormone processing, positive regulation of steroid metabolic process. GSEA also yields the folllowing 22 GO terms related to response to stimulus, protein processing: cellular protein containing complex assembly, cellular response to (CRT) oxygen containing compound, negative regulation of response to stimulus, response to biotic stimulus, response to growth factor, response to endogenous stimulus, response to oxygen containing compound, response to extracellular stimulus, regulation of response to external stimulus negative regulation of response to external stimulus, protein complex oligomerization, protein phosphorylation, regulation of defense response regulation of protein modification process, positive regulation of protein modification process regulation of response to stress, positive regulation of protein metabolic process, response to chemokine, positive regulation of defense response, protein homooligomerazation, multicellular organismal response to stress.

| Designation | Availability | Sample ID | Sample description |
|---|---|---|---|
| Control | Yes | GSM4847382 | A1: MCF7_WT_control |
| Control | Yes | GSM4847383 | A2: MCF7_WT_estrogen_4h |
| Heat Shock | Yes | GSM4847384 | A3: MCF7_WT_HS_1h/2h_rec |
| Control | Yes | GSM4847385 | B1: MCF7_SCR_control |
| Control | Yes | GSM4847387 | B2: MCF7_SCR_estrogen_4h |
| Heat Shock | Yes | GSM4847388 | B3: MCF7_SCR_HS_1h/2h_rec |
| exclude | Yes | GSM4847389 | C1: MCF7_shHSF1_control |
| exclude | Yes | GSM4847390 | C2: MCF7_shHSF1_estrogen_4h |
| exclude | Yes | GSM4847392 | C3: MCF7_shHSF1_HS_1h/2h_rec |
| Control | Yes | GSM4847393 | D1: MCF7_MIX_control |
| Control | Yes | GSM4847394 | D2: MCF7_MIX_estrogen_4h |
| Heat Shock | Yes | GSM4847395 | D3: MCF7_MIX_HS_1h/2h_rec |
| exclude | Yes | GSM4847397 | E1: MCF7_KO#1_control |
| exclude | Yes | GSM4847398 | E2: MCF7_KO#1_estrogen_4h |
| exclude | Yes | GSM4847399 | E3: MCF7_KO#1_HS_1h/2h_rec |
| exclude | Yes | GSM4847400 | F1: MCF7_KO#2_control |
| exclude | Yes | GSM4847402 | F2: MCF7_KO#2_estrogen_4h |
| exclude | Yes | GSM4847403 | F3: MCF7_KO#2_HS_1h/2h_rec |

In GSE164834, the transcriptomic profile of SW620 cancer cells was assessed via mRNAseq analysis following heat-shock both with and without silencing of the Lonp1 gene. Lonp1 was thought to localize only to the mitochondrial matrix and exhibit chaperone functions. The study detected that as much as 22% of Lonp1 localization was also nuclear in response to heat-stress. Paper results excerpt: “We first analyzed the changes in the gene expression caused by HS. In this condition, 442 genes were upregulated and 148 downregulated in cells treated with siLonp1 when compared to cells kept at 37 °C. In cells treated with a scramble siRNA, HS determined the upregulation of 376 genes and the downregulation of 47 genes (Fig. 4A). Genes upregulated during HS in Lonp1-silenced cells were largely overlapping with those upregulated in cells treated with a scramble siRNA, but 125 of them (25%) are uniquely upregulated when Lonp1 was silenced, indicating that silencing of Lonp1 modified HS response (Fig. 4B). As expected, gene ontology analysis revealed that, when compared to control samples, genes upregulated in HS were significantly enriched in “response to unfolded protein”, “protein refolding” and “regulation of cellular response to heat” biological processes, either in cells treated with a Lonp1 siRNA or with a scrambled, control siRNA (Supplementary Table 1)”

| Designation | Availability | Sample ID | Sample description |
|---|---|---|---|
| Control | Yes | GSM5020463 | Ctrl_siCtrl_rep1 |
| Control | Yes | GSM5020464 | Ctrl_siCtrl_rep2 |
| Control | Yes | GSM5020465 | Ctrl_siCtrl_rep3 |
| Control | exclude | GSM5020466 | Ctrl_siLONP1_rep1 |
| Control | exclude | GSM5020467 | Ctrl_siLONP1_rep2 |
| Control | exclude | GSM5020468 | Ctrl_siLONP1_rep3 |
| HeatShock | Yes | GSM5020469 | HS_siCtrl_rep1 |
| HeatShock | Yes | GSM5020470 | HS_siCtrl_rep2 |
| HeatShock | Yes | GSM5020471 | HS_siCtrl_rep3 |
| HeatShock | exclude | GSM5020472 | HS_siLONP1_rep1 |
| HeatShock | exclude | GSM5020473 | HS_siLONP1_rep2 |
| HeatShock | exclude | GSM5020474 | HS_siLONP1_rep3 |
| HeatShock | Yes | GSM5020475 | HS_REC_siCtrl_rep1 |
| HeatShock | Yes | GSM5020476 | HS_REC_siCtrl_rep2 |
| HeatShock | Yes | GSM5020477 | HS_REC_siCtrl_rep3 |
| HeatShock | exclude | GSM5020478 | HS_REC_siLONP1_rep1 |

| | | | |
|---|---|---|---|
| HeatShock | exclude | GSM5020479 | HS_REC_siLONP1_rep2 |
| HeatShock | exclude | GSM5020480 | HS_REC_siLONP1_rep3 |

In GSE182960, heat stress was used to induce and validate differential alternative splicing patterns in HCT116RmAC cells. Heat stressed samples were designated by explicit heat-shocked labeling in GSM identifiers. Control samples were designated as any sample not explicitly heat-stressed. Paper results excerpt: We examined by RNA-seq the effect on gene expression and alternative splicing of cohesin and BRD4 deletion alone or together under physiological and heat shock conditions. Targeted degradation of RAD21 was mediated by an auxin-inducible degron (AID) system (HCT116RmAC cells, RAD21 alleles are fused with a minimal AID tag) and the targeted depletion of BRD4 in HCT116RmAC cells was achieved by treating with MZ1, a proteolysis targeted chimera (PROTAC), under conditions leading to selective degradation of BRD4.

| Designation | available | Sample ID | Sample description |
|---|---|---|---|
| Control | yes | GSM5543727 | 1_RP1_DMSO |
| Control | yes | GSM5543728 | 2_RP2_DMSO |
| Control | yes | GSM5543729 | 3_R3_DMSO |
| Control | yes | GSM5543730 | 4_RP1_MZ1 |
| Control | yes | GSM5543731 | 5_RP2_MZ1 |
| Control | yes | GSM5543732 | 6_R3_MZ1 |
| Control | yes | GSM5543733 | 7_RP1_MZ1_plus_IAA |
| Control | NO | GSM5543734 | 8_RP2_MZ1_plus_IAA |
| Control | yes | GSM5543735 | 9_R3_MZ1_plus_IAA |
| Control | yes | GSM5543736 | 10_RP1_PBS |
| Control | yes | GSM5543737 | 11_RP2_PBS |
| Control | yes | GSM5543738 | 12_R3_PBS |
| Control | yes | GSM5543739 | 13_RP1_IAA |
| Control | yes | GSM5543740 | 14_RP2_IAA |
| Control | yes | GSM5543741 | 15_R3_IAA |

| | | | |
|---|---|---|---|
| Control | yes | GSM5543742 | 1_DMSONHSR1 |
| Control | yes | GSM5543743 | 2_DMSONHSR2 |
| Control | yes | GSM5543744 | 3_DMSONHSR3 |
| Control | yes | GSM5543745 | 4_MZ1NHSR1 |
| Control | yes | GSM5543746 | 5_MZ1NHSR2 |
| Control | yes | GSM5543747 | 6_MZ1NHSR3 |
| Control | yes | GSM5543748 | 7_IAANHSR1 |
| Control | yes | GSM5543749 | 8_IAANHSR2 |
| Control | yes | GSM5543750 | 9_IAANHSR3 |
| Control | yes | GSM5543751 | 10_MZ1_IAANHSR1 |
| Control | yes | GSM5543752 | 11_MZ1_IAANHSR2 |
| Control | yes | GSM5543753 | 12_MZ1_IAANHSR3 |
| Heat-Shock | yes | GSM5543754 | 13_DMSOHSR1 |
| Heat-Shock | yes | GSM5543755 | 14_DMSOHSR2 |
| Heat-Shock | yes | GSM5543756 | 15_DMSOHSR3 |
| Heat-Shock | yes | GSM5543757 | 16_MZ1HSR1 |
| Heat-Shock | yes | GSM5543758 | 17_MZ1HSR2 |
| Heat-Shock | yes | GSM5543759 | 18_MZ1HSR3 |
| Heat-Shock | yes | GSM5543760 | 19_IAAHSR1 |
| Heat-Shock | yes | GSM5543761 | 20_IAAHSR2 |
| Heat-Shock | yes | GSM5543762 | 21_IAAHSR3 |
| Heat-Shock | yes | GSM5543763 | 22_MZ1_IAAHSR1 |
| Heat-Shock | yes | GSM5543764 | 23_MZ1_IAAHSR2 |

| | | | |
|---|---|---|---|
| Heat-Shock | yes | GSM5543765 | 24_MZ1_IAAHSR3 |
