## Supplementary material for "PRRGO: A Tool for Visualizing and Mapping Globally Expressed Genes in Public Gene Expression Omnibus RNA-Sequencing Studies to PageRank-scored Gene Ontology Terms": S2_GEO2R-PRRGO_Complete_Workflow

### GEO2R to PRRGO Complete Workflow

1. Navigate to [https://www.ncbi.nlm.nih.gov/geo/geo2r/](https://www.ncbi.nlm.nih.gov/geo/geo2r/) on a web browser.
2. Enter the GSE ID (GSE######) of interest in the text input next to "GEO accession" and press "Set."

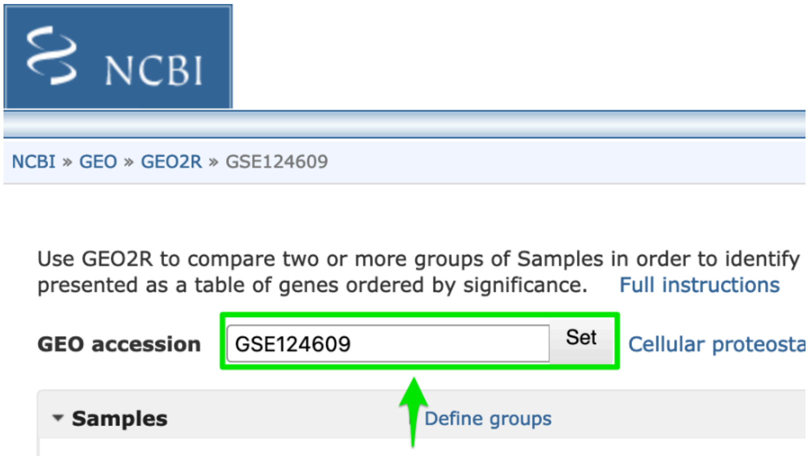


3. Click "Define Groups."

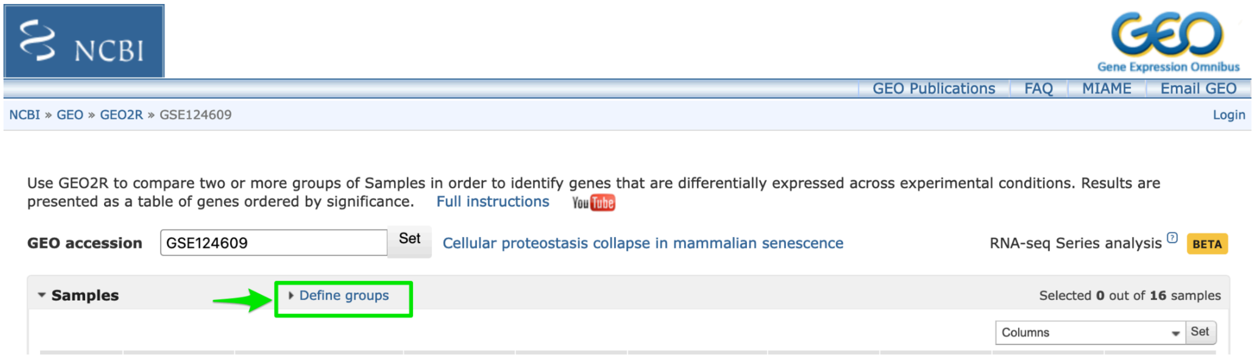


4. In the text input below "Enter a group name: " type "Heat Shock" (or other treatment group) and then "Control" pressing Enter after each. Ensure that "Control" is displayed below Heat Shock for the correct comparison calculations.

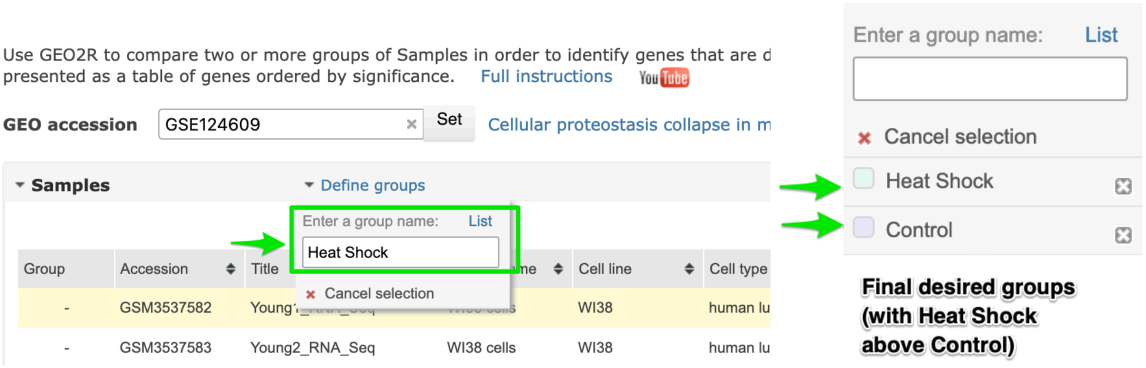


5. Select the samples for the treatment/Heat Shock group by control-clicking or shift-clicking the relevant rows. Control-clicking selects multiple individual rows, while shift-clicking selects a range of rows. Then, click Heat Shock.

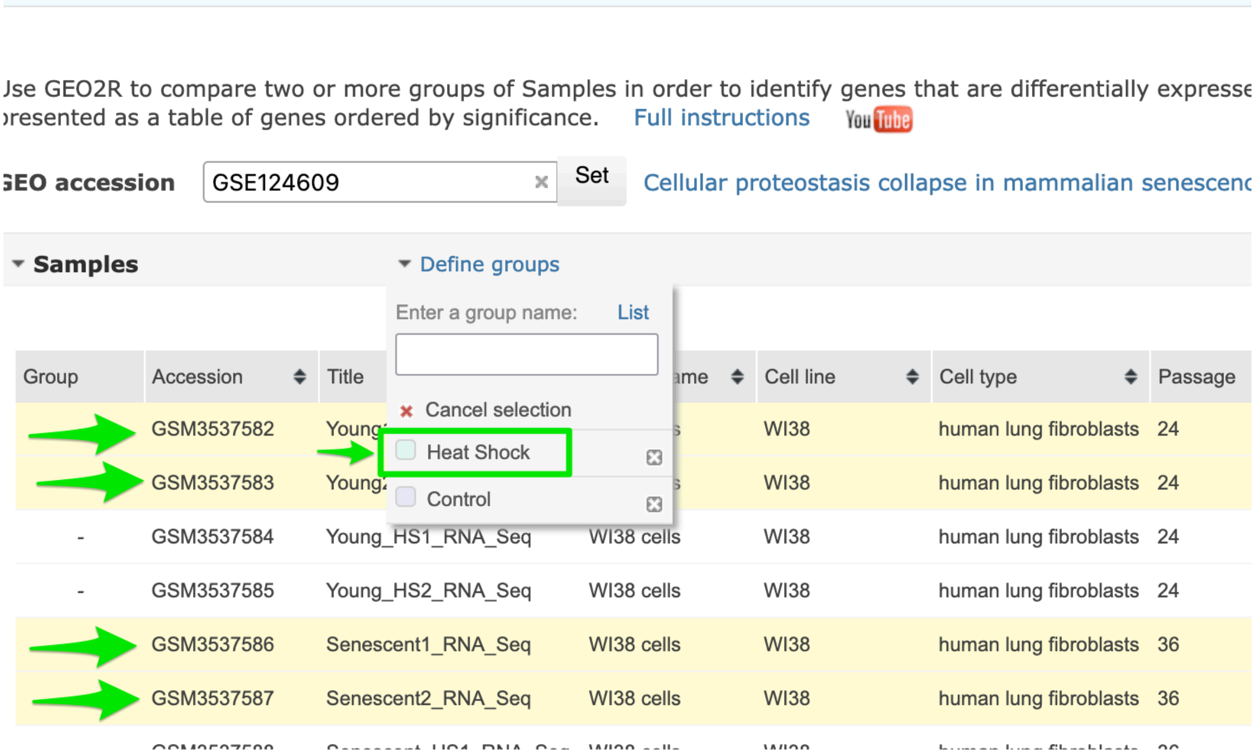

6. Verify that the correct rows are selected in the treatment/Heat Shock group based on the color and Group column.

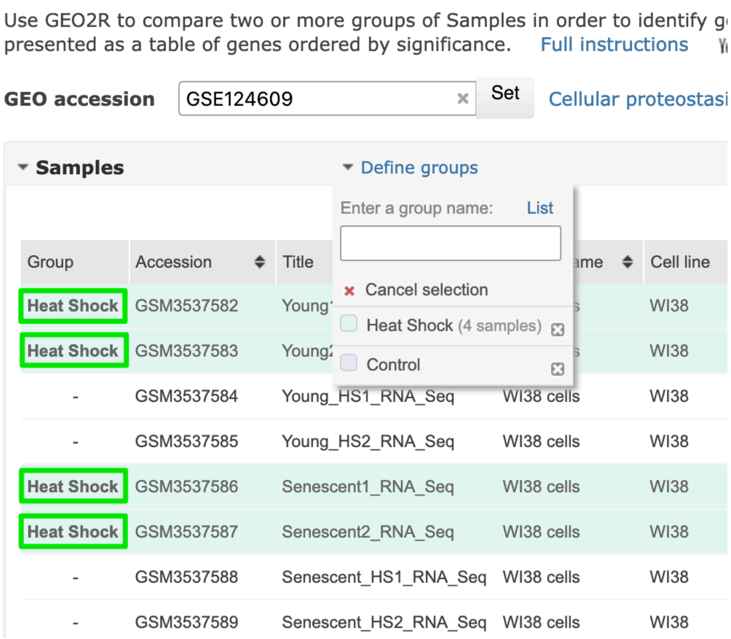


7. Repeat steps 5 and 6 for the Control group. The selected samples for each group are shown below for the example GSE.

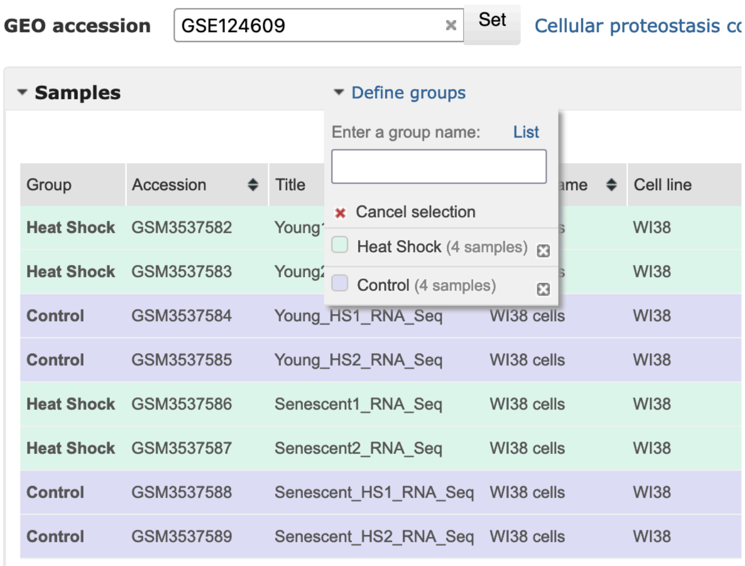

8. Scroll down to the GEO2R tab. Click the Analyze button.

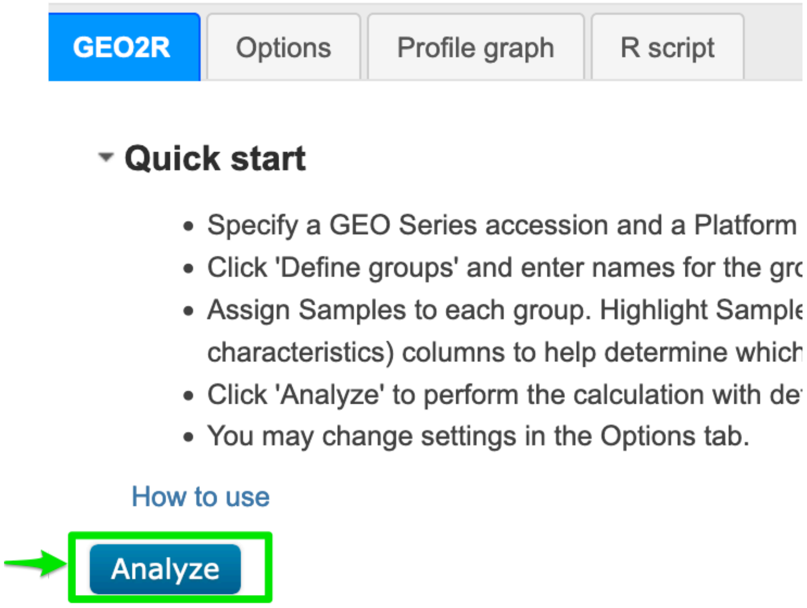


9. Once analysis is complete (may take a minute), click “Download full table” under “Top differentially expressed genes.” The table should download as a tsv file.

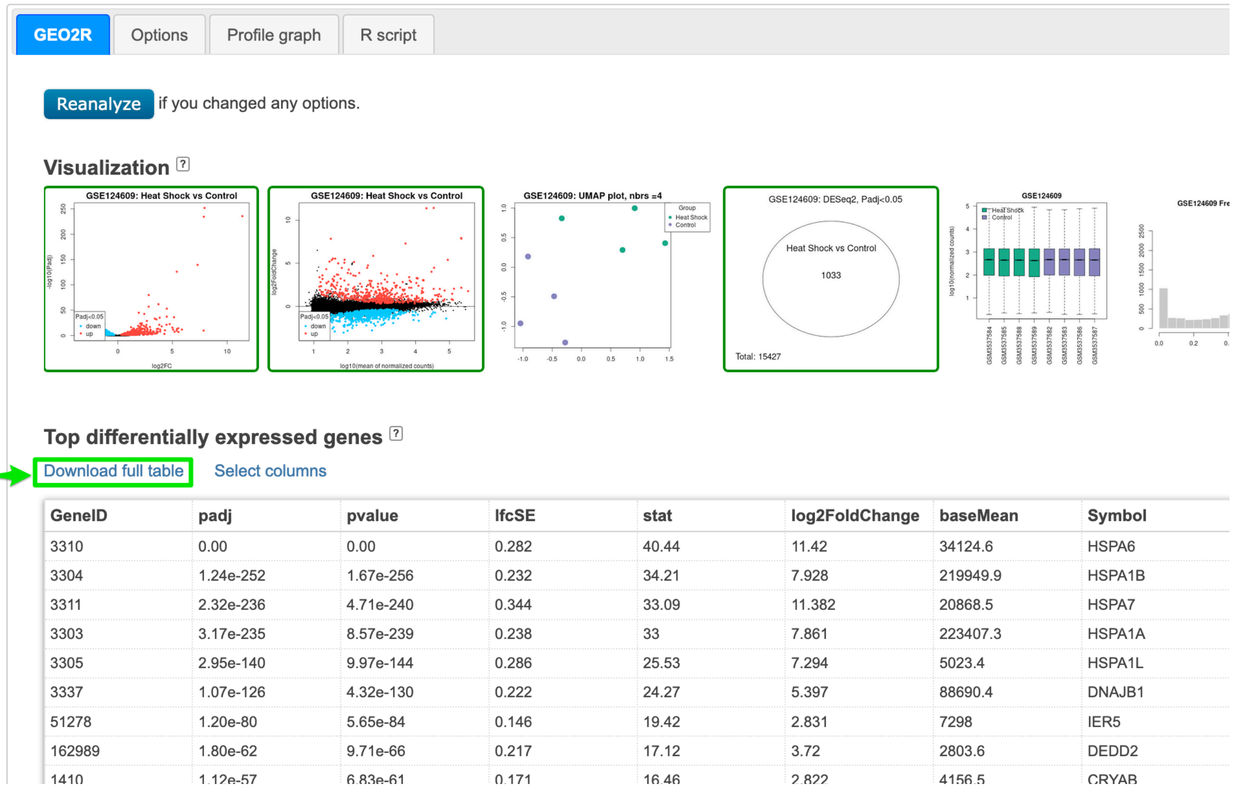


10. (Follow the installation instructions on the PRRGO Github README.md if you have not completed these yet.) Open a shell (Terminal, Powershell, etc.) and

change the working directory to the PRRGO project folder. Run python manage.py runserver. Open a web browser page with the following url: http://127.0.0.1:8000/go-viz.

11. Input the keyword of interest and number of Gene Ontology (GO) terms to filter. Click on "Browse" and click on one or more GEO2R tsv files. Control-click (Windows) or Command-click (Mac) to select multiple files. Press Submit.

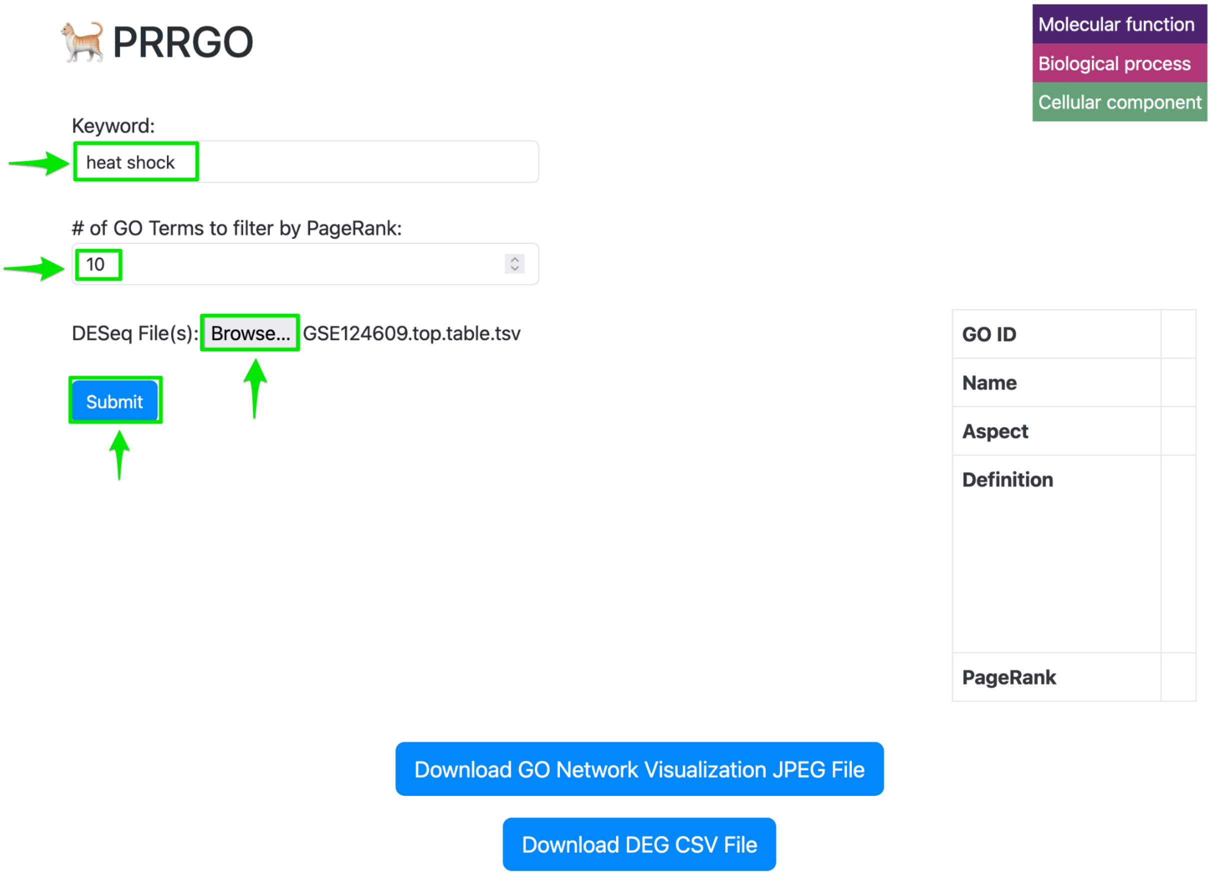


12. After about 15-30 seconds the GO network visualization should appear. Click and drag on the background to pan. Scroll to zoom in/out in the visualization. Click on specific GO term nodes of interest to display relevant GO term information and the table of all differentially expressed genes (DEG) that map to that GO term at the bottom of the page. At this step, you can also click the blue buttons at the bottom to export the visualization as a JPEG or the complete GO

term to DEG mapping as a csv file.

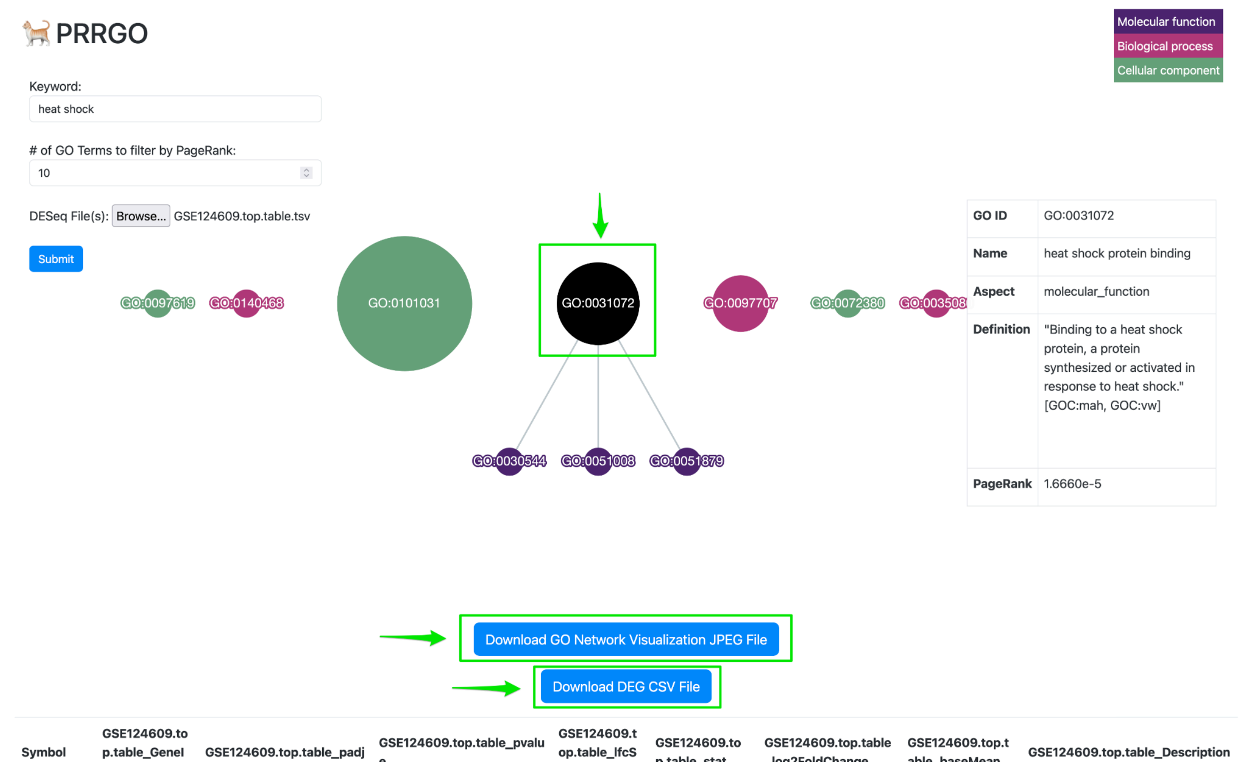


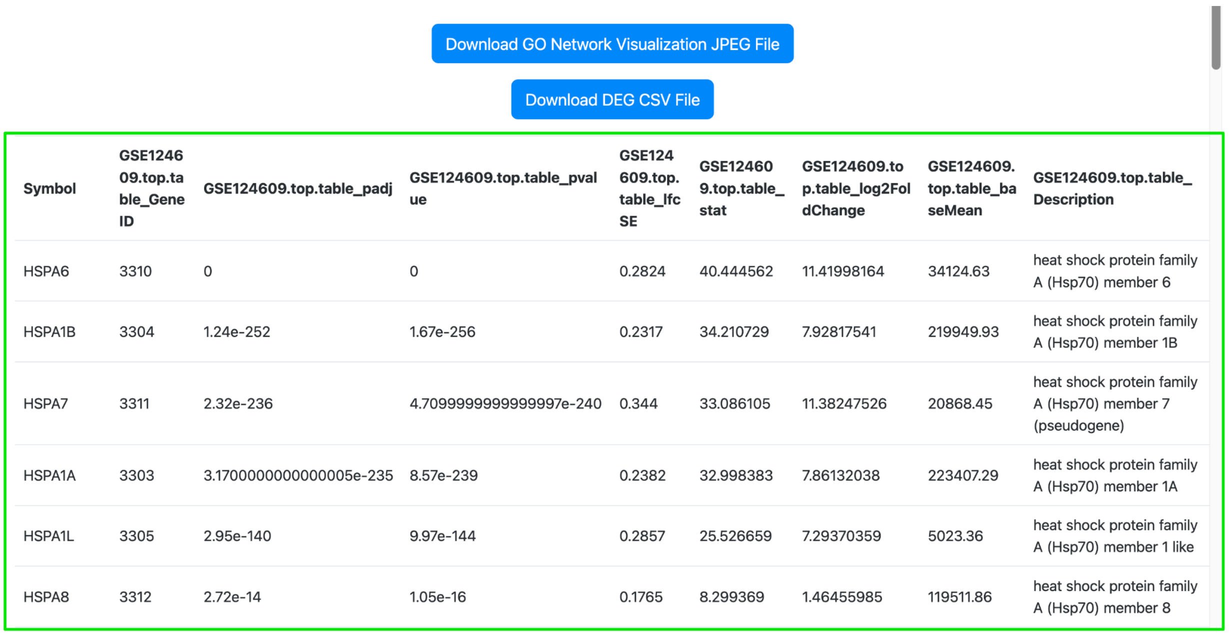


| Symbol | GSE124609.top.table_GeneID | GSE124609.top.table_padj | GSE124609.top.table_pvalue | GSE124609.top.table_lfcSE | GSE124609.top.table_stat | GSE124609.top.table_log2FoldChange | GSE124609.top.table_baseMean | GSE124609.top.table_Description |
|---|---|---|---|---|---|---|---|---|
| HSPA6 | 3310 | 0 | 0 | 0.2824 | 40.444562 | 11.41998164 | 34124.63 | heat shock protein family A (Hsp70) member 6 |
| HSPA1B | 3304 | 1.24e-252 | 1.67e-256 | 0.2317 | 34.210729 | 7.92817541 | 219949.93 | heat shock protein family A (Hsp70) member 1B |
| HSPA7 | 3311 | 2.32e-236 | 4.7099999999999997e-240 | 0.344 | 33.086105 | 11.38247526 | 20868.45 | heat shock protein family A (Hsp70) member 7 (pseudogene) |
| HSPA1A | 3303 | 3.1700000000000005e-235 | 8.57e-239 | 0.2382 | 32.998383 | 7.86132038 | 223407.29 | heat shock protein family A (Hsp70) member 1A |
| HSPA1L | 3305 | 2.95e-140 | 9.97e-144 | 0.2857 | 25.526659 | 7.29370359 | 5023.36 | heat shock protein family A (Hsp70) member 1 like |
| HSPA8 | 3312 | 2.72e-14 | 1.05e-16 | 0.1765 | 8.299369 | 1.46455985 | 119511.86 | heat shock protein family A (Hsp70) member 8 |

*Scroll down after clicking on GO term for DEG table*

13. Close the server with Control-c in the shell (Terminal or other).
