## Supplementary material for "PRRGO: A Tool for Visualizing and Mapping Globally Expressed Genes in Public Gene Expression Omnibus RNA-Sequencing Studies to PageRank-scored Gene Ontology Terms": S4_DEGMetaAnalysis: S4_DEGMetaAnalysis.html

 DGE Meta-Analysis of Nine Gene Expression Omnibus GSE Studies 


Code 

- Show All Code
- Hide All Code

### DGE Meta-Analysis of Nine Gene Expression Omnibus GSE Studies

###### Luis Solano

```
#import libraries
library(tidyverse)
library(UpSetR)
```

```
#import data
GSE124609<-read.csv2('C:/Users/lesolano/Documents/Prrgo_DropboxDL/PRRGO/Validation Studies/GSE124609/GSE124609.ControlvsHS.top.table.tsv', sep = '\t', header = TRUE)
GSE123980<-read.csv2('C:/Users/lesolano/Documents/Prrgo_DropboxDL/PRRGO/Validation Studies/GSE123980/GSE123980.ControlvsHS.top.table.tsv', sep = '\t', header = TRUE)
GSE73471<-read.csv2('C:/Users/lesolano/Documents/Prrgo_DropboxDL/PRRGO/Validation Studies/GSE73471/GSE73471.HSvsControl.top.table.tsv', sep = '\t', header = TRUE)
GSE164834<-read.csv2('C:/Users/lesolano/Documents/Prrgo_DropboxDL/PRRGO/Validation Studies/GSE164834/GSE164834.ControlvsHeatshock.top.table.tsv', sep = '\t', header = TRUE)
GSE66448<-read.csv2('C:/Users/lesolano/Documents/Prrgo_DropboxDL/PRRGO/Validation Studies/GSE66448/GSE66448.ControlvsHS.top.table.tsv', sep = '\t', header = TRUE)
GSE124510<-read.csv2('C:/Users/lesolano/Documents/Prrgo_DropboxDL/PRRGO/Validation Studies/GSE124510/GSE124510.HeatshockvsControl.top.table.tsv', sep = '\t', header = TRUE)
GSE132447<-read.csv2('C:/Users/lesolano/Documents/Prrgo_DropboxDL/PRRGO/Validation Studies/GSE132447/GSE132447.ControlvsHeatshock.top.table.tsv', sep = '\t', header = TRUE)
GSE159802<-read.csv2('C:/Users/lesolano/Documents/Prrgo_DropboxDL/PRRGO/Validation Studies/GSE159802/GSE159802.ControlvsHeatShock.top.table.tsv', sep = '\t', header = TRUE)
GSE182960<-read.csv2('C:/Users/lesolano/Documents/Prrgo_DropboxDL/PRRGO/Validation Studies/GSE182960/GSE182960.HeatshockvsControl.top.table.tsv', sep = '\t', header = TRUE)

#formatting
columns_to_convert <- c('padj', 'log2FoldChange')
GSE124609[columns_to_convert] <- lapply(GSE124609[columns_to_convert], as.numeric)
GSE123980[columns_to_convert] <- lapply(GSE123980[columns_to_convert], as.numeric)
GSE73471[columns_to_convert] <- lapply(GSE73471[columns_to_convert], as.numeric)
GSE164834[columns_to_convert] <- lapply(GSE164834[columns_to_convert], as.numeric)
GSE66448[columns_to_convert] <- lapply(GSE66448[columns_to_convert], as.numeric)
GSE124510[columns_to_convert] <- lapply(GSE124510[columns_to_convert], as.numeric)
GSE132447[columns_to_convert] <- lapply(GSE132447[columns_to_convert], as.numeric)
GSE159802[columns_to_convert] <- lapply(GSE159802[columns_to_convert], as.numeric)
GSE182960[columns_to_convert] <- lapply(GSE182960[columns_to_convert], as.numeric)
```

### 1 Data Wrangle

```
#GSE124609
GSE124609.Detected<-GSE124609$Symbol

GSE124609.Up.PC.sig <-GSE124609 %>%
  filter(grepl("protein-coding", GeneType)) %>%
  filter(padj < 0.05) %>%
  filter(log2FoldChange < 0) %>%
   select(Symbol) %>% pull() 

GSE124609.Up.PC.sigLT1 <-GSE124609 %>%
  filter(grepl("protein-coding", GeneType)) %>%
  filter(padj < 0.05) %>%
  filter(log2FoldChange < -1) %>%
   select(Symbol) %>% pull() 

GSE124609.Down.PC.sig <-GSE124609 %>%
  filter(grepl("protein-coding", GeneType)) %>%
  filter(padj < 0.05) %>%
  filter(log2FoldChange > 0) %>%
   select(Symbol) %>% pull()

GSE124609.Down.PC.sigLT1 <-GSE124609 %>%
  filter(grepl("protein-coding", GeneType)) %>%
  filter(padj < 0.05) %>%
  filter(log2FoldChange > 1) %>%
   select(Symbol) %>% pull()

GSE124609_GOMF_Detected <- GSE124609$GOFunctionID %>%
  strsplit("///") %>%
  unlist() %>%
  unique()

GSE124609_GOMF.PC.sig <- GSE124609 %>%
  filter(grepl("protein-coding", GeneType)) %>%
  filter(padj < 0.05) %>%
  pull(GOFunctionID) %>%
  strsplit("///") %>%
  unlist() %>%
  unique()

GSE124609_GOBP_Detected <- GSE124609$GOProcessID %>%
  strsplit("///") %>%
  unlist() %>%
  unique()

GSE124609_GOBP.PC.sig <- GSE124609 %>%
  filter(grepl("protein-coding", GeneType)) %>%
  filter(padj < 0.05) %>%
  pull(GOProcessID) %>%
  strsplit("///") %>%
  unlist() %>%
  unique()

GSE124609_GOCC_Detected <- GSE124609$GOComponentID %>%
  strsplit("///") %>%
  unlist() %>%
  unique()

GSE124609_GOCC.PC.sig <- GSE124609 %>%
  filter(grepl("protein-coding", GeneType)) %>%
  filter(padj < 0.05) %>%
  pull(GOComponentID) %>%
  strsplit("///") %>%
  unlist() %>%
  unique()

#GSE123980
GSE123980.Detected<-GSE123980$Symbol

GSE123980.Up.PC.sig <-GSE123980 %>%
  filter(grepl("protein-coding", GeneType)) %>%
  filter(padj < 0.05) %>%
  filter(log2FoldChange < 0) %>%
   select(Symbol) %>% pull() 

GSE123980.Up.PC.sigLT1 <-GSE123980 %>%
  filter(grepl("protein-coding", GeneType)) %>%
  filter(padj < 0.05) %>%
  filter(log2FoldChange < -1) %>%
   select(Symbol) %>% pull() 

GSE123980.Down.PC.sig <-GSE123980 %>%
  filter(grepl("protein-coding", GeneType)) %>%
  filter(padj < 0.05) %>%
  filter(log2FoldChange > 0) %>%
   select(Symbol) %>% pull()

GSE123980.Down.PC.sigLT1 <-GSE123980 %>%
  filter(grepl("protein-coding", GeneType)) %>%
  filter(padj < 0.05) %>%
  filter(log2FoldChange > 1) %>%
   select(Symbol) %>% pull()

GSE123980_GOMF_Detected <- GSE123980$GOFunctionID %>%
  strsplit("///") %>%
  unlist() %>%
  unique()

GSE123980_GOMF.PC.sig <- GSE123980 %>%
  filter(grepl("protein-coding", GeneType)) %>%
  filter(padj < 0.05) %>%
  pull(GOFunctionID) %>%
  strsplit("///") %>%
  unlist() %>%
  unique()

GSE123980_GOBP_Detected <- GSE123980$GOProcessID %>%
  strsplit("///") %>%
  unlist() %>%
  unique()

GSE123980_GOBP.PC.sig <- GSE123980 %>%
  filter(grepl("protein-coding", GeneType)) %>%
  filter(padj < 0.05) %>%
  pull(GOProcessID) %>%
  strsplit("///") %>%
  unlist() %>%
  unique()

GSE123980_GOCC_Detected <- GSE123980$GOComponentID %>%
  strsplit("///") %>%
  unlist() %>%
  unique()

GSE123980_GOCC.PC.sig <- GSE123980 %>%
  filter(grepl("protein-coding", GeneType)) %>%
  filter(padj < 0.05) %>%
  pull(GOComponentID) %>%
  strsplit("///") %>%
  unlist() %>%
  unique()

#GSE73471
GSE73471.Detected<-GSE73471$Symbol

GSE73471.Up.PC.sig <-GSE73471 %>%
  filter(grepl("protein-coding", GeneType)) %>%
  filter(padj < 0.05) %>%
  filter(log2FoldChange > 0) %>%
   select(Symbol) %>% pull() 

GSE73471.Up.PC.sigLT1 <-GSE73471 %>%
  filter(grepl("protein-coding", GeneType)) %>%
  filter(padj < 0.05) %>%
  filter(log2FoldChange > 1) %>%
   select(Symbol) %>% pull() 

GSE73471.Down.PC.sig <-GSE73471 %>%
  filter(grepl("protein-coding", GeneType)) %>%
  filter(padj < 0.05) %>%
  filter(log2FoldChange < 0) %>%
   select(Symbol) %>% pull()

GSE73471.Down.PC.sigLT1 <-GSE73471 %>%
  filter(grepl("protein-coding", GeneType)) %>%
  filter(padj < 0.05) %>%
  filter(log2FoldChange < -1) %>%
   select(Symbol) %>% pull()

GSE73471_GOMF_Detected <- GSE73471$GOFunctionID %>%
  strsplit("///") %>%
  unlist() %>%
  unique()

GSE73471_GOMF.PC.sig <- GSE73471 %>%
  filter(grepl("protein-coding", GeneType)) %>%
  filter(padj < 0.05) %>%
  pull(GOFunctionID) %>%
  strsplit("///") %>%
  unlist() %>%
  unique()

GSE73471_GOBP_Detected <- GSE73471$GOProcessID %>%
  strsplit("///") %>%
  unlist() %>%
  unique()

GSE73471_GOBP.PC.sig <- GSE73471 %>%
  filter(grepl("protein-coding", GeneType)) %>%
  filter(padj < 0.05) %>%
  pull(GOProcessID) %>%
  strsplit("///") %>%
  unlist() %>%
  unique()

GSE73471_GOCC_Detected <- GSE73471$GOComponentID %>%
  strsplit("///") %>%
  unlist() %>%
  unique()

GSE73471_GOCC.PC.sig <- GSE73471 %>%
  filter(grepl("protein-coding", GeneType)) %>%
  filter(padj < 0.05) %>%
  pull(GOComponentID) %>%
  strsplit("///") %>%
  unlist() %>%
  unique()

#GSE164834
GSE164834.Detected<-GSE164834$Symbol

GSE164834.Up.PC.sig <-GSE164834 %>%
  filter(grepl("protein-coding", GeneType)) %>%
  filter(padj < 0.05) %>%
  filter(log2FoldChange < 0) %>%
   select(Symbol) %>% pull() 

GSE164834.Up.PC.sigLT1 <-GSE164834 %>%
  filter(grepl("protein-coding", GeneType)) %>%
  filter(padj < 0.05) %>%
  filter(log2FoldChange < -1) %>%
   select(Symbol) %>% pull()

GSE164834.Down.PC.sig <-GSE164834 %>%
  filter(grepl("protein-coding", GeneType)) %>%
  filter(padj < 0.05) %>%
  filter(log2FoldChange > 0) %>%
   select(Symbol) %>% pull()

GSE164834.Down.PC.sigLT1 <-GSE164834 %>%
  filter(grepl("protein-coding", GeneType)) %>%
  filter(padj < 0.05) %>%
  filter(log2FoldChange > 1) %>%
   select(Symbol) %>% pull()

GSE164834_GOMF_Detected <- GSE164834$GOFunctionID %>%
  strsplit("///") %>%
  unlist() %>%
  unique()

GSE164834_GOMF.PC.sig <- GSE164834 %>%
  filter(grepl("protein-coding", GeneType)) %>%
  filter(padj < 0.05) %>%
  pull(GOFunctionID) %>%
  strsplit("///") %>%
  unlist() %>%
  unique()

GSE164834_GOBP_Detected <- GSE164834$GOProcessID %>%
  strsplit("///") %>%
  unlist() %>%
  unique()

GSE164834_GOBP.PC.sig <- GSE164834 %>%
  filter(grepl("protein-coding", GeneType)) %>%
  filter(padj < 0.05) %>%
  pull(GOProcessID) %>%
  strsplit("///") %>%
  unlist() %>%
  unique()

GSE164834_GOCC_Detected <- GSE164834$GOComponentID %>%
  strsplit("///") %>%
  unlist() %>%
  unique()

GSE164834_GOCC.PC.sig <- GSE164834 %>%
  filter(grepl("protein-coding", GeneType)) %>%
  filter(padj < 0.05) %>%
  pull(GOComponentID) %>%
  strsplit("///") %>%
  unlist() %>%
  unique()

#GSE66448
GSE66448.Detected<-GSE66448$Symbol

GSE66448.Up.PC.sig <-GSE66448 %>%
  filter(grepl("protein-coding", GeneType)) %>%
  filter(padj < 0.05) %>%
  filter(log2FoldChange < 0) %>%
   select(Symbol) %>% pull() 

GSE66448.Up.PC.sigLT1 <-GSE66448 %>%
  filter(grepl("protein-coding", GeneType)) %>%
  filter(padj < 0.05) %>%
  filter(log2FoldChange < -1) %>%
   select(Symbol) %>% pull()

GSE66448.Down.PC.sig <-GSE66448 %>%
  filter(grepl("protein-coding", GeneType)) %>%
  filter(padj < 0.05) %>%
  filter(log2FoldChange > 0) %>%
   select(Symbol) %>% pull()

GSE66448.Down.PC.sigLT1 <-GSE66448 %>%
  filter(grepl("protein-coding", GeneType)) %>%
  filter(padj < 0.05) %>%
  filter(log2FoldChange > 1) %>%
   select(Symbol) %>% pull()

GSE66448_GOMF_Detected <- GSE66448$GOFunctionID %>%
  strsplit("///") %>%
  unlist() %>%
  unique()

GSE66448_GOMF.PC.sig <- GSE66448 %>%
  filter(grepl("protein-coding", GeneType)) %>%
  filter(padj < 0.05) %>%
  pull(GOFunctionID) %>%
  strsplit("///") %>%
  unlist() %>%
  unique()

GSE66448_GOBP_Detected <- GSE66448$GOProcessID %>%
  strsplit("///") %>%
  unlist() %>%
  unique()

GSE66448_GOBP.PC.sig <- GSE66448 %>%
  filter(grepl("protein-coding", GeneType)) %>%
  filter(padj < 0.05) %>%
  pull(GOProcessID) %>%
  strsplit("///") %>%
  unlist() %>%
  unique()

GSE66448_GOCC_Detected <- GSE66448$GOComponentID %>%
  strsplit("///") %>%
  unlist() %>%
  unique()

GSE66448_GOCC.PC.sig <- GSE66448 %>%
  filter(grepl("protein-coding", GeneType)) %>%
  filter(padj < 0.05) %>%
  pull(GOComponentID) %>%
  strsplit("///") %>%
  unlist() %>%
  unique()

#GSE124510
GSE124510.Detected<-GSE124510$Symbol

GSE124510.Up.PC.sig <-GSE124510 %>%
  filter(grepl("protein-coding", GeneType)) %>%
  filter(padj < 0.05) %>%
  filter(log2FoldChange > 0) %>%
   select(Symbol) %>% pull() 

GSE124510.Up.PC.sigLT1 <-GSE124510 %>%
  filter(grepl("protein-coding", GeneType)) %>%
  filter(padj < 0.05) %>%
  filter(log2FoldChange > 1) %>%
   select(Symbol) %>% pull()

GSE124510.Down.PC.sig <-GSE124510 %>%
  filter(grepl("protein-coding", GeneType)) %>%
  filter(padj < 0.05) %>%
  filter(log2FoldChange < 0) %>%
   select(Symbol) %>% pull()

GSE124510.Down.PC.sigLT1 <-GSE124510 %>%
  filter(grepl("protein-coding", GeneType)) %>%
  filter(padj < 0.05) %>%
  filter(log2FoldChange < -1) %>%
   select(Symbol) %>% pull()

GSE124510_GOMF_Detected <- GSE124510$GOFunctionID %>%
  strsplit("///") %>%
  unlist() %>%
  unique()

GSE124510_GOMF.PC.sig <- GSE124510 %>%
  filter(grepl("protein-coding", GeneType)) %>%
  filter(padj < 0.05) %>%
  pull(GOFunctionID) %>%
  strsplit("///") %>%
  unlist() %>%
  unique()

GSE124510_GOBP_Detected <- GSE124510$GOProcessID %>%
  strsplit("///") %>%
  unlist() %>%
  unique()

GSE124510_GOBP.PC.sig <- GSE124510 %>%
  filter(grepl("protein-coding", GeneType)) %>%
  filter(padj < 0.05) %>%
  pull(GOProcessID) %>%
  strsplit("///") %>%
  unlist() %>%
  unique()

GSE124510_GOCC_Detected <- GSE124510$GOComponentID %>%
  strsplit("///") %>%
  unlist() %>%
  unique()

GSE124510_GOCC.PC.sig <- GSE124510 %>%
  filter(grepl("protein-coding", GeneType)) %>%
  filter(padj < 0.05) %>%
  pull(GOComponentID) %>%
  strsplit("///") %>%
  unlist() %>%
  unique()

#GSE132447
GSE132447.Detected<-GSE132447$Symbol

GSE132447.Up.PC.sig <-GSE132447 %>%
  filter(grepl("protein-coding", GeneType)) %>%
  filter(padj < 0.05) %>%
  filter(log2FoldChange < 0) %>%
   select(Symbol) %>% pull() 

GSE132447.Up.PC.sigLT1 <-GSE132447 %>%
  filter(grepl("protein-coding", GeneType)) %>%
  filter(padj < 0.05) %>%
  filter(log2FoldChange < -1) %>%
   select(Symbol) %>% pull()

GSE132447.Down.PC.sig <-GSE132447 %>%
  filter(grepl("protein-coding", GeneType)) %>%
  filter(padj < 0.05) %>%
  filter(log2FoldChange > 0) %>%
   select(Symbol) %>% pull()

GSE132447.Down.PC.sigLT1 <-GSE132447 %>%
  filter(grepl("protein-coding", GeneType)) %>%
  filter(padj < 0.05) %>%
  filter(log2FoldChange > 1) %>%
   select(Symbol) %>% pull()

GSE132447_GOMF_Detected <- GSE132447$GOFunctionID %>%
  strsplit("///") %>%
  unlist() %>%
  unique()

GSE132447_GOMF.PC.sig <- GSE132447 %>%
  filter(grepl("protein-coding", GeneType)) %>%
  filter(padj < 0.05) %>%
  pull(GOFunctionID) %>%
  strsplit("///") %>%
  unlist() %>%
  unique()

GSE132447_GOBP_Detected <- GSE132447$GOProcessID %>%
  strsplit("///") %>%
  unlist() %>%
  unique()

GSE132447_GOBP.PC.sig <- GSE132447 %>%
  filter(grepl("protein-coding", GeneType)) %>%
  filter(padj < 0.05) %>%
  pull(GOProcessID) %>%
  strsplit("///") %>%
  unlist() %>%
  unique()

GSE132447_GOCC_Detected <- GSE132447$GOComponentID %>%
  strsplit("///") %>%
  unlist() %>%
  unique()

GSE132447_GOCC.PC.sig <- GSE132447 %>%
  filter(grepl("protein-coding", GeneType)) %>%
  filter(padj < 0.05) %>%
  pull(GOComponentID) %>%
  strsplit("///") %>%
  unlist() %>%
  unique()

#GSE159802
GSE159802.Detected<-GSE159802$Symbol

GSE159802.Up.PC.sig <-GSE159802 %>%
  filter(grepl("protein-coding", GeneType)) %>%
  filter(padj < 0.05) %>%
  filter(log2FoldChange < 0) %>%
   select(Symbol) %>% pull() 

GSE159802.Up.PC.sigLT1 <-GSE159802 %>%
  filter(grepl("protein-coding", GeneType)) %>%
  filter(padj < 0.05) %>%
  filter(log2FoldChange < -1) %>%
   select(Symbol) %>% pull()

GSE159802.Down.PC.sig <-GSE159802 %>%
  filter(grepl("protein-coding", GeneType)) %>%
  filter(padj < 0.05) %>%
  filter(log2FoldChange > 0) %>%
   select(Symbol) %>% pull()

GSE159802.Down.PC.sigLT1 <-GSE159802 %>%
  filter(grepl("protein-coding", GeneType)) %>%
  filter(padj < 0.05) %>%
  filter(log2FoldChange > 1) %>%
   select(Symbol) %>% pull()

GSE159802_GOMF_Detected <- GSE159802$GOFunctionID %>%
  strsplit("///") %>%
  unlist() %>%
  unique()

GSE159802_GOMF.PC.sig <- GSE159802 %>%
  filter(grepl("protein-coding", GeneType)) %>%
  filter(padj < 0.05) %>%
  pull(GOFunctionID) %>%
  strsplit("///") %>%
  unlist() %>%
  unique()

GSE159802_GOBP_Detected <- GSE159802$GOProcessID %>%
  strsplit("///") %>%
  unlist() %>%
  unique()

GSE159802_GOBP.PC.sig <- GSE159802 %>%
  filter(grepl("protein-coding", GeneType)) %>%
  filter(padj < 0.05) %>%
  pull(GOProcessID) %>%
  strsplit("///") %>%
  unlist() %>%
  unique()

GSE159802_GOCC_Detected <- GSE159802$GOComponentID %>%
  strsplit("///") %>%
  unlist() %>%
  unique()

GSE159802_GOCC.PC.sig <- GSE159802 %>%
  filter(grepl("protein-coding", GeneType)) %>%
  filter(padj < 0.05) %>%
  pull(GOComponentID) %>%
  strsplit("///") %>%
  unlist() %>%
  unique()

#GSE182960
GSE182960.Detected<-GSE182960$Symbol

GSE182960.Up.PC.sig <-GSE182960 %>%
  filter(grepl("protein-coding", GeneType)) %>%
  filter(padj < 0.05) %>%
  filter(log2FoldChange > 0) %>%
   select(Symbol) %>% pull() 

GSE182960.Up.PC.sigLT1 <-GSE182960 %>%
  filter(grepl("protein-coding", GeneType)) %>%
  filter(padj < 0.05) %>%
  filter(log2FoldChange > 1) %>%
   select(Symbol) %>% pull()

GSE182960.Down.PC.sig <-GSE182960 %>%
  filter(grepl("protein-coding", GeneType)) %>%
  filter(padj < 0.05) %>%
  filter(log2FoldChange < 0) %>%
   select(Symbol) %>% pull()

GSE182960.Down.PC.sigLT1 <-GSE182960 %>%
  filter(grepl("protein-coding", GeneType)) %>%
  filter(padj < 0.05) %>%
  filter(log2FoldChange < -1) %>%
   select(Symbol) %>% pull()

GSE182960_GOMF_Detected <- GSE182960$GOFunctionID %>%
  strsplit("///") %>%
  unlist() %>%
  unique()

GSE182960_GOMF.PC.sig <- GSE182960 %>%
  filter(grepl("protein-coding", GeneType)) %>%
  filter(padj < 0.05) %>%
  pull(GOFunctionID) %>%
  strsplit("///") %>%
  unlist() %>%
  unique()

GSE182960_GOBP_Detected <- GSE182960$GOProcessID %>%
  strsplit("///") %>%
  unlist() %>%
  unique()

GSE182960_GOBP.PC.sig <- GSE182960 %>%
  filter(grepl("protein-coding", GeneType)) %>%
  filter(padj < 0.05) %>%
  pull(GOProcessID) %>%
  strsplit("///") %>%
  unlist() %>%
  unique()

GSE182960_GOCC_Detected <- GSE182960$GOComponentID %>%
  strsplit("///") %>%
  unlist() %>%
  unique()

GSE182960_GOCC.PC.sig <- GSE182960 %>%
  filter(grepl("protein-coding", GeneType)) %>%
  filter(padj < 0.05) %>%
  pull(GOComponentID) %>%
  strsplit("///") %>%
  unlist() %>%
  unique()
```

### 2 Upset plots

#### 2.1 DEG Upset Plots

##### 2.1.1 Up Genes

```
Upset_UpHS_input <- list(`GSE124609.Up.PC.sig` = GSE124609.Up.PC.sig,
                   `GSE123980.Up.PC.sig` = GSE123980.Up.PC.sig,
                   `GSE73471.Up.PC.sig` = GSE73471.Up.PC.sig,
                   `GSE164834.Up.PC.sig` = GSE164834.Up.PC.sig,
                   `GSE66448.Up.PC.sig` = GSE66448.Up.PC.sig,
                   `GSE124510.Up.PC.sig` = GSE124510.Up.PC.sig,
                   `GSE132447.Up.PC.sig` = GSE132447.Up.PC.sig,
                   `GSE159802.Up.PC.sig` = GSE159802.Up.PC.sig,
                   `GSE182960.Up.PC.sig` = GSE182960.Up.PC.sig)

Upset_UpHS<-upset(fromList(Upset_UpHS_input), nsets = 9, order.by = "freq")
Upset_UpHS
```

##### 2.1.2 Down Genes

```
Upset_DownHS_input <- list(`GSE124609.Down.PC.sig` = GSE124609.Down.PC.sig,
                   `GSE123980.Down.PC.sig` = GSE123980.Down.PC.sig,
                   `GSE73471.Down.PC.sig` = GSE73471.Down.PC.sig,
                   `GSE164834.Down.PC.sig` = GSE164834.Down.PC.sig,
                   `GSE66448.Down.PC.sig` = GSE66448.Down.PC.sig,
                   `GSE124510.Down.PC.sig` = GSE124510.Down.PC.sig,
                   `GSE132447.Down.PC.sig` = GSE132447.Down.PC.sig,
                   `GSE159802.Down.PC.sig` = GSE159802.Down.PC.sig,
                   `GSE182960.Down.PC.sig` = GSE182960.Down.PC.sig)

Upset_DownHS<-upset(fromList(Upset_DownHS_input), nsets = 9, order.by = "freq")
Upset_DownHS
```

##### 2.1.3 Total DEG Intersection

```
Upset_TotalHS_input <- list(`GSE124609.Up.PC.sig` = GSE124609.Up.PC.sig,
                   `GSE123980.Up.PC.sig` = GSE123980.Up.PC.sig,
                   `GSE73471.Up.PC.sig` = GSE73471.Up.PC.sig,
                   `GSE164834.Up.PC.sig` = GSE164834.Up.PC.sig,
                   `GSE66448.Up.PC.sig` = GSE66448.Up.PC.sig,
                   `GSE124510.Up.PC.sig` = GSE124510.Up.PC.sig,
                   `GSE132447.Up.PC.sig` = GSE132447.Up.PC.sig,
                   `GSE159802.Up.PC.sig` = GSE159802.Up.PC.sig,
                   `GSE182960.Up.PC.sig` = GSE182960.Up.PC.sig,
                   `GSE124609.Down.PC.sig` = GSE124609.Down.PC.sig,
                   `GSE123980.Down.PC.sig` = GSE123980.Down.PC.sig,
                   `GSE73471.Down.PC.sig` = GSE73471.Down.PC.sig,
                   `GSE164834.Down.PC.sig` = GSE164834.Down.PC.sig,
                   `GSE66448.Down.PC.sig` = GSE66448.Down.PC.sig,
                   `GSE124510.Down.PC.sig` = GSE124510.Down.PC.sig,
                   `GSE132447.Down.PC.sig` = GSE132447.Down.PC.sig,
                   `GSE159802.Down.PC.sig` = GSE159802.Down.PC.sig,
                   `GSE182960.Down.PC.sig` = GSE182960.Down.PC.sig)

Upset_TotalHS<-upset(fromList(Upset_TotalHS_input), nsets = 9, order.by = "freq")
Upset_TotalHS
```

#### 2.2 GO Upset Plots

##### 2.2.1 GO Molecular Function Intersection

```
Upset_GOMF_input <- list(`GSE124609_GOMF.PC.sig` = GSE124609_GOMF.PC.sig,
                   `GSE123980_GOMF.PC.sig` = GSE123980_GOMF.PC.sig,
                   `GSE73471_GOMF.PC.sig` = GSE73471_GOMF.PC.sig,
                   `GSE164834_GOMF.PC.sig` = GSE164834_GOMF.PC.sig,
                   `GSE66448_GOMF.PC.sig` = GSE66448_GOMF.PC.sig,
                   `GSE124510_GOMF.PC.sig` = GSE124510_GOMF.PC.sig,
                   `GSE132447_GOMF.PC.sig` = GSE132447_GOMF.PC.sig,
                   `GSE159802_GOMF.PC.sig` = GSE159802_GOMF.PC.sig,
                   `GSE182960_GOMF.PC.sig` = GSE182960_GOMF.PC.sig)

Upset_GOMF<-upset(fromList(Upset_GOMF_input), nsets = 9, order.by = "freq")
Upset_GOMF
```

##### 2.2.2 GO Biological Process Intersection

```
Upset_GOBP_input <- list(`GSE124609_GOBP.PC.sig` = GSE124609_GOBP.PC.sig,
                   `GSE123980_GOBP.PC.sig` = GSE123980_GOBP.PC.sig,
                   `GSE73471_GOBP.PC.sig` = GSE73471_GOBP.PC.sig,
                   `GSE164834_GOBP.PC.sig` = GSE164834_GOBP.PC.sig,
                   `GSE66448_GOBP.PC.sig` = GSE66448_GOBP.PC.sig,
                   `GSE124510_GOBP.PC.sig` = GSE124510_GOBP.PC.sig,
                   `GSE132447_GOBP.PC.sig` = GSE132447_GOBP.PC.sig,
                   `GSE159802_GOBP.PC.sig` = GSE159802_GOBP.PC.sig,
                   `GSE182960_GOBP.PC.sig` = GSE182960_GOBP.PC.sig)

Upset_GOBP<-upset(fromList(Upset_GOBP_input), nsets = 9, order.by = "freq")
Upset_GOBP
```

##### 2.2.3 GO Cellular Component Intersection

```
Upset_GOCC_input <- list(`GSE124609_GOCC.PC.sig` = GSE124609_GOCC.PC.sig,
                   `GSE123980_GOCC.PC.sig` = GSE123980_GOCC.PC.sig,
                   `GSE73471_GOCC.PC.sig` = GSE73471_GOCC.PC.sig,
                   `GSE164834_GOCC.PC.sig` = GSE164834_GOCC.PC.sig,
                   `GSE66448_GOCC.PC.sig` = GSE66448_GOCC.PC.sig,
                   `GSE124510_GOCC.PC.sig` = GSE124510_GOCC.PC.sig,
                   `GSE132447_GOCC.PC.sig` = GSE132447_GOCC.PC.sig,
                   `GSE159802_GOCC.PC.sig` = GSE159802_GOCC.PC.sig,
                   `GSE182960_GOCC.PC.sig` = GSE182960_GOCC.PC.sig)

Upset_GOCC<-upset(fromList(Upset_GOCC_input), nsets = 9, order.by = "freq")
Upset_GOCC
```

#### 2.3 HSR GO term Hits

```
HSGO <- list('GO:0101031', 'GO:0031072', 'GO:0051008',  'GO:0051879', 'GO:0030544', 'GO:0140468', 'GO:0072380')

find_matching_entries <- function(input_list) {
  m_entries <- input_list[input_list %in% HSGO]
  return(m_entries)
}

#GSE124609
GSE124609_GOterms <- list(GSE124609_GOMF.PC.sig, GSE124609_GOBP.PC.sig, GSE124609_GOCC.PC.sig)

# New list to store matching entries for each list
GSE124609matchingGOterms <- list()

# Loop through each list in GSE124609_GOterms
for (i in seq_along(GSE124609_GOterms)) {
  matching_entries <- find_matching_entries(GSE124609_GOterms[[i]])
  GSE124609matchingGOterms <- c(GSE124609matchingGOterms, matching_entries)
}

#GSE123980
GSE123980_GOterms <- list(GSE123980_GOMF.PC.sig, GSE123980_GOBP.PC.sig, GSE123980_GOCC.PC.sig)

# New list to store matching entries for each list
GSE123980matchingGOterms <- list()

# Loop through each list in GSE123980_GOterms
for (i in seq_along(GSE123980_GOterms)) {
  matching_entries <- find_matching_entries(GSE123980_GOterms[[i]])
  GSE123980matchingGOterms <- c(GSE123980matchingGOterms, matching_entries)
}

#GSE73471
GSE73471_GOterms <- list(GSE73471_GOMF.PC.sig, GSE73471_GOBP.PC.sig, GSE73471_GOCC.PC.sig)

# New list to store matching entries for each list
GSE73471matchingGOterms <- list()

# Loop through each list in GSE73471_GOterms
for (i in seq_along(GSE73471_GOterms)) {
  matching_entries <- find_matching_entries(GSE73471_GOterms[[i]])
  GSE73471matchingGOterms <- c(GSE73471matchingGOterms, matching_entries)
}

#GSE164834
GSE164834_GOterms <- list(GSE164834_GOMF.PC.sig, GSE164834_GOBP.PC.sig, GSE164834_GOCC.PC.sig)

# New list to store matching entries for each list
GSE164834matchingGOterms <- list()

# Loop through each list in GSE164834_GOterms
for (i in seq_along(GSE164834_GOterms)) {
  matching_entries <- find_matching_entries(GSE164834_GOterms[[i]])
  GSE164834matchingGOterms <- c(GSE164834matchingGOterms, matching_entries)
}

#GSE66448
GSE66448_GOterms <- list(GSE66448_GOMF.PC.sig, GSE66448_GOBP.PC.sig, GSE66448_GOCC.PC.sig)

# New list to store matching entries for each list
GSE66448matchingGOterms <- list()

# Loop through each list in GSE66448_GOterms
for (i in seq_along(GSE66448_GOterms)) {
  matching_entries <- find_matching_entries(GSE66448_GOterms[[i]])
  GSE66448matchingGOterms <- c(GSE66448matchingGOterms, matching_entries)
}

#GSE124510
GSE124510_GOterms <- list(GSE124510_GOMF.PC.sig, GSE124510_GOBP.PC.sig, GSE124510_GOCC.PC.sig)

# New list to store matching entries for each list
GSE124510matchingGOterms <- list()

# Loop through each list in GSE124510_GOterms
for (i in seq_along(GSE124510_GOterms)) {
  matching_entries <- find_matching_entries(GSE124510_GOterms[[i]])
  GSE124510matchingGOterms <- c(GSE124510matchingGOterms, matching_entries)
}

#GSE132447
GSE132447_GOterms <- list(GSE132447_GOMF.PC.sig, GSE132447_GOBP.PC.sig, GSE132447_GOCC.PC.sig)

# New list to store matching entries for each list
GSE132447matchingGOterms <- list()

# Loop through each list in GSE132447_GOterms
for (i in seq_along(GSE132447_GOterms)) {
  matching_entries <- find_matching_entries(GSE132447_GOterms[[i]])
  GSE132447matchingGOterms <- c(GSE132447matchingGOterms, matching_entries)
}

#GSE159802
GSE159802_GOterms <- list(GSE159802_GOMF.PC.sig, GSE159802_GOBP.PC.sig, GSE159802_GOCC.PC.sig)

# New list to store matching entries for each list
GSE159802matchingGOterms <- list()

# Loop through each list in GSE159802_GOterms
for (i in seq_along(GSE159802_GOterms)) {
  matching_entries <- find_matching_entries(GSE159802_GOterms[[i]])
  GSE159802matchingGOterms <- c(GSE159802matchingGOterms, matching_entries)
}

#GSE182960
GSE182960_GOterms <- list(GSE182960_GOMF.PC.sig, GSE182960_GOBP.PC.sig, GSE182960_GOCC.PC.sig)

# New list to store matching entries for each list
GSE182960matchingGOterms <- list()

# Loop through each list in GSE182960_GOterms
for (i in seq_along(GSE182960_GOterms)) {
  matching_entries <- find_matching_entries(GSE182960_GOterms[[i]])
  GSE182960matchingGOterms <- c(GSE182960matchingGOterms, matching_entries)
}


Upset_HSGO_input<- list(`GSE124609 GO Terms: 0031072, 0030544, 0051879, 0140468, 0101031` = GSE124609matchingGOterms,
`GSE123980 GO Terms: 0030544, 0031072, 0051879, 0140468, 0101031, 0072380` = GSE123980matchingGOterms,
`GSE73471 GO Terms: 0031072, 0030544, 0051879, 0140468, 0101031` = GSE73471matchingGOterms,
`GSE164834 GO Terms: 0031072,0030544,0051879,0101031` = GSE164834matchingGOterms,
`GSE66448 GO Terms: 0031072, 0030544, 0051879, 0140468, 0101031` = GSE66448matchingGOterms,
`GSE124510 GO Terms: 0030544, 0051879, 0031072, 0140468, 0101031, 0072380` = GSE124510matchingGOterms,
`GSE132447 GO Terms: 0031072, 0030544, 0051879, 0140468, 0101031` = GSE132447matchingGOterms,
`GSE159802 GO Terms: 0031072, 0030544, 0051879, 0051008, 0101031` = GSE159802matchingGOterms,
`GSE182960 GO Terms: 0030544, 0031072, 0051879, 0051008, 0140468, 0101031` = GSE182960matchingGOterms)

Upset_HSGO<-upset(fromList(Upset_HSGO_input), nsets = 9, order.by = "freq")
Upset_HSGO
```

### 3 Descriptive Summary Reporting

#### 3.1 DEGs

GSE124609

Study GSE124609 detected 15427 total genes and 894 statistically
significant (p.adj<0.05) protein coding differentially expressed
genes.

GSE123980

Study GSE123980 detected 22115 total genes and 3691 statistically
significant (p.adj<0.05) protein coding differentially expressed
genes.

GSE73471

Study GSE73471 detected 17416 total genes and 1672 statistically
significant (p.adj<0.05) protein coding differentially expressed
genes.

GSE164834

Study GSE164834 detected 18042 total genes and 4000 statistically
significant (p.adj<0.05) protein coding differentially expressed
genes.

GSE66448

Study GSE66448 detected 17755 total genes and 3140 statistically
significant (p.adj<0.05) protein coding differentially expressed
genes.

GSE124510

Study GSE124510 detected 17896 total genes and 7213 statistically
significant (p.adj<0.05) protein coding differentially expressed
genes.

GSE132447

Study GSE132447 detected 23202 total genes and 2448 statistically
significant (p.adj<0.05) protein coding differentially expressed
genes.

GSE159802

Study GSE159802 detected 16385 total genes and 2521 statistically
significant (p.adj<0.05) protein coding differentially expressed
genes.

GSE182960

Study GSE182960 detected 16773 total genes and 5105 statistically
significant (p.adj<0.05) protein coding differentially expressed
genes.

#### 3.2 Gene Ontology

GSE124609

GEO2R reports significant genes were associated with 970 Gene
Ontology-Molecular Function terms in GSE124609.

GEO2R reports significant genes were associated with 3546 Gene
Ontology-Biological Process terms in GSE124609.

GEO2R reports significant genes were associated with 594 Gene
Ontology-Cellular Component terms in GSE124609.

GSE123980

GEO2R reports significant genes were associated with 2302 Gene
Ontology-Molecular Function terms in GSE123980.

GEO2R reports significant genes were associated with 7202 Gene
Ontology-Biological Process terms in GSE123980.

GEO2R reports significant genes were associated with 1169 Gene
Ontology-Cellular Component terms in GSE123980.

GSE73471

GEO2R reports significant genes were associated with 1483 Gene
Ontology-Molecular Function terms in GSE73471.

GEO2R reports significant genes were associated with 4965 Gene
Ontology-Biological Process terms in GSE73471.

GEO2R reports significant genes were associated with 855 Gene
Ontology-Cellular Component terms in GSE73471.

GSE164834

GEO2R reports significant genes were associated with 2328 Gene
Ontology-Molecular Function terms in GSE164834.

GEO2R reports significant genes were associated with 7381 Gene
Ontology-Biological Process terms in GSE164834.

GEO2R reports significant genes were associated with 1177 Gene
Ontology-Cellular Component terms in GSE164834.

GSE66448

GEO2R reports significant genes were associated with 2014 Gene
Ontology-Molecular Function terms in GSE66448.

GEO2R reports significant genes were associated with 6243 Gene
Ontology-Biological Process terms in GSE66448.

GEO2R reports significant genes were associated with 1093 Gene
Ontology-Cellular Component terms in GSE66448.

GSE124510

GEO2R reports significant genes were associated with 3085 Gene
Ontology-Molecular Function terms in GSE124510.

GEO2R reports significant genes were associated with 9172 Gene
Ontology-Biological Process terms in GSE124510.

GEO2R reports significant genes were associated with 1464 Gene
Ontology-Cellular Component terms in GSE124510.

GSE132447

GEO2R reports significant genes were associated with 1793 Gene
Ontology-Molecular Function terms in GSE132447.

GEO2R reports significant genes were associated with 6126 Gene
Ontology-Biological Process terms in GSE132447.

GEO2R reports significant genes were associated with 918 Gene
Ontology-Cellular Component terms in GSE132447.

GSE159802

GEO2R reports significant genes were associated with 1715 Gene
Ontology-Molecular Function terms in GSE159802.

GEO2R reports significant genes were associated with 5902 Gene
Ontology-Biological Process terms in GSE159802.

GEO2R reports significant genes were associated with 952 Gene
Ontology-Cellular Component terms in GSE159802.

GSE182960

GEO2R reports significant genes were associated with 2598 Gene
Ontology-Molecular Function terms in GSE182960.

GEO2R reports significant genes were associated with 7926 Gene
Ontology-Biological Process terms in GSE182960.

GEO2R reports significant genes were associated with 1299 Gene
Ontology-Cellular Component terms in GSE182960.

### 4 Individual GSE Result Template

X samples from study GSE124609 were processed with GEO2R, which
reports that 15427 total genes had measurable expression (Figure 3A). Of
these detected genes, 342 protein coding genes were calculated as
positively differentially expressed and 552 protein coding genes were
calculated as negatively differentially expressed for a total of 894
protein coding genes at statistically significant (p.adj <0.05)
levels (Figure 3A). Of the statistically significant positively
differentially expressed genes, 181 exhibited log2FC expression >1
and 227 exhibited log2FC expression <-1 for a total of 408 genes with
|log2FC expression|>1 (Figure 3A). Of the statistically significant
differentially expressed genes, GEO2R also reports 970 associated Gene
Ontology-Molecular Function terms, 3546 associated Gene
Ontology-Biological Process terms and 594 associated Gene
Ontology-Cellular Component terms (Figure 3A). These enriched GO terms
contained key HSR indicator GO terms: GO:0031072, GO:0030544, GO:0051879, GO:0140468, GO:0101031.

X samples from study GSE123980 were processed with GEO2R, which
reports that 22115 total genes had measurable expression (Figure 3A). Of
these detected genes, 1714 protein coding genes were calculated as
positively differentially expressed and 1977 protein coding genes were
calculated as negatively differentially expressed for a total of 3691
protein coding genes at statistically significant (p.adj <0.05)
levels (Figure 3A). Of the statistically significant positively
differentially expressed genes, 52 exhibited log2FC expression >1 and
173 exhibited log2FC expression <-1 for a total of 225 genes with
|log2FC expression|>1 (Figure 3A). Of the statistically significant
differentially expressed genes, GEO2R also reports 2302 associated Gene
Ontology-Molecular Function terms, 7202 associated Gene
Ontology-Biological Process terms and 1169 associated Gene
Ontology-Cellular Component terms (Figure 3A). These enriched GO terms
contained key HSR indicator GO terms: GO:0030544, GO:0031072, GO:0051879, GO:0140468, GO:0101031, GO:0072380.

X samples from study GSE73471 were processed with GEO2R, which
reports that 17416 total genes had measurable expression (Figure 3A). Of
these detected genes, 836 protein coding genes were calculated as
positively differentially expressed and 836 protein coding genes were
calculated as negatively differentially expressed for a total of 1672
protein coding genes at statistically significant (p.adj <0.05)
levels (Figure 3A). Of the statistically significant positively
differentially expressed genes, 192 exhibited log2FC expression >1
and 321 exhibited log2FC expression <-1 for a total of 513 genes with
|log2FC expression|>1 (Figure 3A). Of the statistically significant
differentially expressed genes, GEO2R also reports 1483 associated Gene
Ontology-Molecular Function terms, 4965 associated Gene
Ontology-Biological Process terms and 855 associated Gene
Ontology-Cellular Component terms (Figure 3A). These enriched GO terms
contained key HSR indicator GO terms: GO:0031072, GO:0030544, GO:0051879, GO:0140468, GO:0101031.

X samples from study GSE164834 were processed with GEO2R, which
reports that 18042 total genes had measurable expression (Figure 3A). Of
these detected genes, 2147 protein coding genes were calculated as
positively differentially expressed and 1853 protein coding genes were
calculated as negatively differentially expressed for a total of 4000
protein coding genes at statistically significant (p.adj <0.05)
levels (Figure 3A). Of the statistically significant positively
differentially expressed genes, 479 exhibited log2FC expression >1
and 39 exhibited log2FC expression <-1 for a total of 518 genes with
|log2FC expression|>1 (Figure 3A). Of the statistically significant
differentially expressed genes, GEO2R also reports 2328 associated Gene
Ontology-Molecular Function terms, 7381 associated Gene
Ontology-Biological Process terms and 1177 associated Gene
Ontology-Cellular Component terms (Figure 3A). These enriched GO terms
contained key HSR indicator GO terms: GO:0031072, GO:0030544, GO:0051879, GO:0101031.

X samples from study GSE66448 were processed with GEO2R, which
reports that 17755 total genes had measurable expression (Figure 3A). Of
these detected genes, 1547 protein coding genes were calculated as
positively differentially expressed and 1593 protein coding genes were
calculated as negatively differentially expressed for a total of 3140
protein coding genes at statistically significant (p.adj <0.05)
levels (Figure 3A). Of the statistically significant positively
differentially expressed genes, 985 exhibited log2FC expression >1
and 772 exhibited log2FC expression <-1 for a total of 1757 genes
with |log2FC expression|>1 (Figure 3A). Of the statistically
significant differentially expressed genes, GEO2R also reports 2014
associated Gene Ontology-Molecular Function terms, 6243 associated Gene
Ontology-Biological Process terms and 1093 associated Gene
Ontology-Cellular Component terms (Figure 3A). These enriched GO terms
contained key HSR indicator GO terms: GO:0031072, GO:0030544, GO:0051879, GO:0140468, GO:0101031.

X samples from study GSE124510 were processed with GEO2R, which
reports that 17896 total genes had measurable expression (Figure 3A). Of
these detected genes, 3075 protein coding genes were calculated as
positively differentially expressed and 4138 protein coding genes were
calculated as negatively differentially expressed for a total of 7213
protein coding genes at statistically significant (p.adj <0.05)
levels (Figure 3A). Of the statistically significant positively
differentially expressed genes, 1631 exhibited log2FC expression >1
and 2230 exhibited log2FC expression <-1 for a total of 3861 genes
with |log2FC expression|>1 (Figure 3A). Of the statistically
significant differentially expressed genes, GEO2R also reports 3085
associated Gene Ontology-Molecular Function terms, 9172 associated Gene
Ontology-Biological Process terms and 1464 associated Gene
Ontology-Cellular Component terms (Figure 3A). These enriched GO terms
contained key HSR indicator GO terms: GO:0030544, GO:0051879, GO:0031072, GO:0140468, GO:0101031, GO:0072380.

X samples from study GSE132447 were processed with GEO2R, which
reports that 23202 total genes had measurable expression (Figure 3A). Of
these detected genes, 1527 protein coding genes were calculated as
positively differentially expressed and 921 protein coding genes were
calculated as negatively differentially expressed for a total of 2448
protein coding genes at statistically significant (p.adj <0.05)
levels (Figure 3A). Of the statistically significant positively
differentially expressed genes, 977 exhibited log2FC expression >1
and 273 exhibited log2FC expression <-1 for a total of 1250 genes
with |log2FC expression|>1 (Figure 3A). Of the statistically
significant differentially expressed genes, GEO2R also reports 1793
associated Gene Ontology-Molecular Function terms, 6126 associated Gene
Ontology-Biological Process terms and 918 associated Gene
Ontology-Cellular Component terms (Figure 3A). These enriched GO terms
contained key HSR indicator GO terms: GO:0031072, GO:0030544, GO:0051879, GO:0140468, GO:0101031.

X samples from study GSE159802 were processed with GEO2R, which
reports that 16385 total genes had measurable expression (Figure 3A). Of
these detected genes, 1181 protein coding genes were calculated as
positively differentially expressed and 1340 protein coding genes were
calculated as negatively differentially expressed for a total of 2521
protein coding genes at statistically significant (p.adj <0.05)
levels (Figure 3A). Of the statistically significant positively
differentially expressed genes, 672 exhibited log2FC expression >1
and 296 exhibited log2FC expression <-1 for a total of 968 genes with
|log2FC expression|>1 (Figure 3A). Of the statistically significant
differentially expressed genes, GEO2R also reports 1715 associated Gene
Ontology-Molecular Function terms, 5902 associated Gene
Ontology-Biological Process terms and 952 associated Gene
Ontology-Cellular Component terms (Figure 3A). These enriched GO terms
contained key HSR indicator GO terms: GO:0031072, GO:0030544, GO:0051879, GO:0051008, GO:0101031.

X samples from study GSE182960 were processed with GEO2R, which
reports that 16773 total genes had measurable expression (Figure 3A). Of
these detected genes, 2640 protein coding genes were calculated as
positively differentially expressed and 2465 protein coding genes were
calculated as negatively differentially expressed for a total of 5105
protein coding genes at statistically significant (p.adj <0.05)
levels (Figure 3A). Of the statistically significant positively
differentially expressed genes, 361 exhibited log2FC expression >1
and 25 exhibited log2FC expression <-1 for a total of 386 genes with
|log2FC expression|>1 (Figure 3A). Of the statistically significant
differentially expressed genes, GEO2R also reports 2598 associated Gene
Ontology-Molecular Function terms, 7926 associated Gene
Ontology-Biological Process terms and 1299 associated Gene
Ontology-Cellular Component terms (Figure 3A). These enriched GO terms
contained key HSR indicator GO terms: GO:0030544, GO:0031072, GO:0051879, GO:0051008, GO:0140468, GO:0101031.

#### 4.1 Reproducibility

The output from running ‘sessionInfo’ is shown below and details all
packages and version necessary to reproduce the results in this
report.

```
sessionInfo()
```

```
## R version 4.3.1 (2023-06-16 ucrt)
## Platform: x86_64-w64-mingw32/x64 (64-bit)
## Running under: Windows 11 x64 (build 22621)
## 
## Matrix products: default
## 
## 
## locale:
## [1] LC_COLLATE=English_United States.utf8 
## [2] LC_CTYPE=English_United States.utf8   
## [3] LC_MONETARY=English_United States.utf8
## [4] LC_NUMERIC=C                          
## [5] LC_TIME=English_United States.utf8    
## 
## time zone: America/Los_Angeles
## tzcode source: internal
## 
## attached base packages:
## [1] stats     graphics  grDevices utils     datasets  methods   base     
## 
## other attached packages:
##  [1] UpSetR_1.4.0    lubridate_1.9.2 forcats_1.0.0   stringr_1.5.0  
##  [5] dplyr_1.1.3     purrr_1.0.2     readr_2.1.4     tidyr_1.3.0    
##  [9] tibble_3.2.1    ggplot2_3.4.3   tidyverse_2.0.0 knitr_1.43     
## [13] tinytex_0.46    rmarkdown_2.24 
## 
## loaded via a namespace (and not attached):
##  [1] sass_0.4.7        utf8_1.2.3        generics_0.1.3    stringi_1.7.12   
##  [5] hms_1.1.3         digest_0.6.33     magrittr_2.0.3    evaluate_0.21    
##  [9] grid_4.3.1        timechange_0.2.0  fastmap_1.1.1     plyr_1.8.9       
## [13] jsonlite_1.8.7    gridExtra_2.3     fansi_1.0.4       scales_1.2.1     
## [17] codetools_0.2-19  jquerylib_0.1.4   cli_3.6.1         rlang_1.1.1      
## [21] munsell_0.5.0     withr_2.5.0       cachem_1.0.8      yaml_2.3.7       
## [25] tools_4.3.1       tzdb_0.4.0        colorspace_2.1-0  vctrs_0.6.3      
## [29] R6_2.5.1          lifecycle_1.0.3   pkgconfig_2.0.3   pillar_1.9.0     
## [33] bslib_0.5.1       gtable_0.3.4      glue_1.6.2        Rcpp_1.0.11      
## [37] xfun_0.40         tidyselect_1.2.0  highr_0.10        rstudioapi_0.15.0
## [41] farver_2.1.1      htmltools_0.5.6   labeling_0.4.3    compiler_4.3.1
```
